# Developing the Grapevine Hydric Stress Atlas: A Meta-Analysis Resource for Exploring Transcriptome Responses to Drought

**DOI:** 10.1101/2025.03.31.646369

**Authors:** Álvaro Vidal Valenzuela, David Navarro-Payá, Antonio Santiago, Paolo Sonego, Felipe Gainza-Cortés, Mickael Malnoy, José Tomás Matus

**Affiliations:** Research and Innovation centre, Fondazione Edmund Mach, Via Mach 1, 38098 San michelle all’adige(TN), Italy; Institute for Integrative Systems Biology (I2SysBio), Universitat de València-CSIC, Paterna, 46980, Valencia, Spain; Center for Research and Innovation (CII), Viña Concha y Toro, 3550000, Pencahue, Chile; Center Agriculture Food Environment (C3A), University of Trento, via E. Mach 1, 38010 San Michele all’Adige, Italy

**Author notes:** corresponding authorship. first author (Álvaro Vidal Valenzuela).

## Abstract

Climate change poses a significant threat to agriculture, particularly in regions where increased drought periods, abnormal heat, and intensified pest pressure threaten crop productivity. Viticulture, as one of the most economically important crops, is highly vulnerable to these challenges. Understanding the molecular mechanisms underlying drought responses in grapevines is essential for developing innovative molecular breeding strategies aimed at enhancing drought tolerance, improving cultivar resilience, and promoting agricultural sustainability. In this context, transcriptomic meta-analyses have proven valuable for uncovering global regulatory trends, gene co-expression networks and conserved biological responses across diverse conditions. As part of this study, 1107 public transcriptomic datasets from Illumina (631 runs) and ABI SOLID (476 runs) platforms were searched, reclassified and reanalyzed in the latest T2T genome assembly, in order to construct condition, cultivar and tissue-specific gene expression atlases associated with drought stress in grapevine. To facilitate exploration of this data, a web-based application, the Hydric Stress Atlas App (https://plantaeviz.tomsbiolab.com/vitviz/hydric_atlas/), was developed, as part of the Vitis module within the PlantaeViz platform. Together with this tool, we generated a whole-genome co-expression network using the same datasets (https://plantaeviz.tomsbiolab.com/vitviz/networks/non_agg_gcns/T2T/hydric_stress_TI/). This water stress condition-dependent GCN allows to explore and visualize gene co-expression relationships related to stress and identify network hubs holding novel drought stress regulators. We manually curated experimental metadata, and enabled the classification of transcriptomic data by cultivar, tissue, and drought tolerance. Finally, candidate genes associated with drought tolerance were identified via network topology analysis. These genes can be further used as molecular markers, or characterized via gene editing or cisgenesis, providing insights into their molecular roles in drought tolerance. This resource contributes to a deeper understanding of grapevine drought responses, offering a pathway for sustainable viticulture and innovative biotechnological solutions to address climate-related challenges.

## Introduction

The cultivated grapevine (*Vitis vinifera* ssp. *vinifera*, henceforth referred to as *Vitis vinifera*) has shown remarkable adaptability to significant global climatic shifts and human migrations throughout history. It served as a critical energy source for humans since the late Pleistocene, and its domestication during the Holocene coincided with improved climatic conditions and the migration of agrarian communities. Over centuries, the domestication of *Vitis vinifera* paralleled the establishment of sedentary societies and the development of trade networks, particularly during the Bronze Age, facilitating the exchange of elite cultivars across Europe and beyond (Dong *et al*., 2023). Grapes in this period transitioned from a dioecious wild ancestor to an inbreeding monoecious species. This co-evolutionary relationship between grapevine and humanity, spanning several centuries, has deeply shaped its genetic and phenotypic diversity, making it a vital contributor to food and beverage production and an insightful model for understanding human-plant interactions.

Environmental conditions and anthropogenic advancements have profoundly influenced grapevine evolution. In the Anthropocene epoch, human-induced climate change poses an unprecedented challenge, leaving imprints that will persist in geological records for millennia (Roka, 2019). The impacts of climate change on grapevine productivity are evident, intensifying abiotic stressors such as drought, salinity, soil acidification, high temperatures, excessive radiation, and unexpected extreme weather events such as summer storms, late-spring frosts (LSFs), and other meteorological conditions. These stressors not only threaten plant health but also alter pathogen dynamics, leading to the emergence of novel strains and an increased prevalence of diseases like powdery mildew (Singh *et al.,* 2023; Tang *et al.,* 2017). Among abiotic stressors, water deficit, a major contributor to drought conditions, is particularly detrimental. It is responsible for greater annual crop yield losses than the combined effects of all pathogens (Gupta *et al*., 2020).

Drought, characterized by prolonged reductions in precipitation, when accompanied with high temperatures and erratic weather patterns, has profound effects on grapevine physiology, yield, and grape composition (Mishra and Singh, 2010). Drought primarily induces heat and water stress in plants, either concurrently or independently (Jaldhani *et al*., 2022). In grapevines, water deficit affects not only the composition, quality, and yield of grapes but has also resulted in a significant decline in wine production, significantly affecting the wine industry. In 2023, the International Organisation of Vine and Wine (OIV) reported the lowest global wine production in six decades due to adverse climatic conditions, with substantial reductions observed across both hemispheres (OIV, 2023). The economic consequences are severe, threatening an industry valued at USD 483.6 billion in 2023 (IMARC). Furthermore, projections suggest that up to 90% of traditional wine-producing regions in Southern Europe and California could become unsuitable for viticulture by the end of the century due to worsening drought and heat waves (Van Leeuwen *et al*., 2024).

The variability in grapevine responses to water stress is largely influenced by transcriptional changes across *Vitis* species and cultivars (Konecny *et al.,* 2024). These changes affect critical physiological processes such as osmotic adjustment, photosynthesis, phytohormones, and antioxidant activity (Lin *et al*., 2023). Notably, combined water deficit and heat stress conditions elicit more pronounced transcriptional changes than either stress alone, highlighting the importance of studying grapevine responses under realistic environmental scenarios (reviewed Martínez-Luscher *et al*., 2025). To address these challenges, a systems biology approach that interrogates transcriptomic behaviors upon water stress is essential.

Transcriptomic studies have been particularly insightful, addressing key questions related to drought responses, including the role of transcriptional regulation in ABA signaling, nitrogen metabolism, and secondary metabolism pathways (Cerda and Alvarez, 2023; Wong *et al*., 2016; Orduña *et al*., 2022). Moreover, transcriptomic studies have identified splicing variants linked to stress adaptation and revealed stress-specific gene expression patterns, such as those regulating raffinose synthesis in drought-stressed grapevines (Trenti *et al*., 2021). Despite these advances, there is still room for improvement. Transcriptomic meta-analyses seek to combine various independent studies to elucidate common phenomic traits and have shown remarkable potential for gene discovery in the context of biotic stress resistance. This is exemplified by the identification of *VviMLO* genes in grapevine, first described as a family (Feechan *et al*., 2008) and later transcriptionally linked to powdery mildew infection (Winterhagen *et al*., 2008). These genes were eventually silenced using RNAi and confirmed as susceptibility genes (S-genes, Pessina *et al*., 2016). Building on this work, as well as studies in other species for S-gene knockout, the Arabidopsis susceptibility genes *AtDLO* and At*DMR6* were analyzed. Through phylogenetic and transcriptomic analysis, the *Vitis VviDMR6-1* gene was identified as an *S*-gene for Downy mildew infection (Pirrello *et al*., 2022), and was subsequently edited using CRISPR (Giacomelli et al., 2022; Giacomelli *et al*., 2023; Djennane *et al*., 2023). Consequently, mining public transcriptomic data has become a standard practice to achieve desired phenotypes.

In this work, we reanalyzed public grape RNA-seq data, for which we catalogued all available runs present in the Sequence Read Archive (SRA) of NCBI, selecting those related to water stress, using Illumina or ABI-SOLID sequencing platforms. These were manually curated to be selected based on the experimental design described on each associated publication. Using the filtered experiments, the Hydric Stress Atlas (https://plantaeviz.tomsbiolab.com/vitviz/hydric_atlas/) and the non-aggregated gene co-expression network (https://plantaeviz.tomsbiolab.com/vitviz/networks/non_agg_gcns/T2T/hydric_stress_TI/) were constructed. This application offers different tools that help explore gene expression under drought conditions, where the effects on expression can be studied under different intensities of stress, four different tissues, cultivars, and rootstocks catalogued as either sensitive or tolerant. To demonstrate its utility, genes associated with the synthesis and signaling pathways of Abscisic Acid (ABA) under water stress conditions were analyzed to identify canonical response genes. Additionally, pathways involved in cellular reprogramming in response to stress were studied, such as the modification of stomatal density through the disruption of the cellular differentiation mechanism at early developmental stages of guard cells, as the protodermal phase. Finally, co-expression networks involved in the response to water stress were examined to identify potential transcription factors that regulate gene expression under stress conditions. The development of such comprehensive resources provides valuable insights into the molecular mechanisms underlying drought tolerance and can guide future strategies for grapevine improvement.

## Results

### Exploration of drought experiments in grapevine transcriptomic public data

This study analyzed drought-responsive transcriptomic data in *Vitis* species, selecting experiments based on their design and normalizing results according to keyword-based categorization. To identify relevant RNA-seq datasets, a search was conducted in the SRA database of NCBI in December 2024, using the following query in the title and/or abstract: “(grapevine [Title/Abstract] OR grape [Title/Abstract] OR Vitis [Title/Abstract] OR V. vinifera [Title/Abstract]) AND (drought OR water stress OR water deficiency OR water deficit stress) AND (transcriptome [Title/Abstract] OR transcript profiling [Title/Abstract] OR mRNA expression [Title/Abstract] OR RNA-Seq [Title/Abstract] OR RNASeq [Title/Abstract] OR RNA Sequencing [Title/Abstract] OR RNA-Sequencing [Title/Abstract])”. This search retrieved 1,386 runs, which were then filtered to include only experiments meeting statistical significance criteria, resulting in 997 runs. These datasets encompass samples from roots, leaves, shoots, and berries, with genetic backgrounds including cvs. ‘Riparia Gloire’, ‘Cabernet Sauvignon’, ‘Ramsey’, ‘101-14’, ‘M4’, ‘Chardonnay’, ‘Merlot’, ‘Pinot Noir’, ‘Semillon’, ‘Cabernet Volos’, ‘Tocai Friulano’, ‘Riesling’, and an unnamed *Vitis vinifera x Vitis girdiana* hybrid. The final selection includes 10 studies that applied different levels of water stress, capturing diverse responses across conditions.

The ten identified studies stand out for their unique experimental designs, particularly those involving time-course experiments or comparisons between tolerant (anisohydric) and sensitive (isohydric) cultivars. Notably, the bioproject PRJNA516950 by Cochetel et al. (2020), examined drought responses in the wine cultivar *Vitis vinifera* cv. ‘Cabernet Sauvignon’ clone 8 (CS), along with the wild species *Vitis riparia* (RG), *Vitis champinii* (RM), and a *Vitis vinifera* x *girdiana* hybrid (SC). Plants were subjected to water stress based on relative soil water content (RSWC) at field capacity, with natural evaporation occurring over 7 and 14 days. RNA was extracted from at least three biological replicates per treatment, while control plants were maintained at 100% RSWC. During the 7-day drought period, RSWC declined to 50%, leading to a stationary phase of stem water potential, which dropped exponentially from -0.5 MPa across all genotypes.

Several studies evaluated the transcriptomic differences between rootstocks. In the bioproject PRJNA429560 by Kahdka *et al*. (2019), *V. riparia* Michx. were subjected to 7- and 14-day water stress treatments with three replicates each. Shoot elongation and physiological parameters were monitored, revealing changes in primary shoot length and node number between 7 and 10. Stem water potential and net photosynthesis rate significantly decreased by day 7. For bioprojects PRJNA226228 and PRJNA226229 (Vitulo *et al*., 2014; Meggio *et al*., 2014; Corso *et al.,* 2015), two drought-tolerant rootstock genotypes, 101.14 Millardet et de Grasset (*V. riparia* x *V. rupestris*) and a new ([(*V. vinifera × V. berlandier*i) × *V. berlandieri* cv. Resseguier n. 1], named M4), were studied. Water stress was induced by progressively reducing RSWC to 30% for stressed plants and 80% for controls, with measurements taken at 0 (T0), 2 (T1), 4 (T2), 7(T3), and 10 (T4) days. From day 6, RSWC in stressed plants remained constant at ∼30%.

A few studies have explored tolerance among *Vitis vinifera* cultivars. In the bioproject PRJNA662522 by Tan *et al*. (2023), *V. vinifera* L. cv. ‘Cabernet Sauvignon’ was subjected to drought stress defined based on stomatal conductance (gs) to water vapor. Stress was defined as gs values between 75 and 100 mmol/m²/s, with plants maintained under drought conditions for 10 days, distinguishing moderate from severe stress. The bioproject PRJEB44212 focused on cv. ‘Fleurtai’ and ‘Cabernet Volos’, which experienced field drought over two years. Drought severity was determined using the average midday stem water potential. In the bioproject PRJNA268857 by Ghan *et al*. (2015), berries from five different *Vitis vinifera* cultivars (‘Cabernet Sauvignon’, ‘Merlot’, ‘Pinot Noir’, ‘Chardonnay’, and ‘Semillon’), underwent field drought treatments, with midday stem water potential maintained at ∼−0.8 MPa for stressed vines and ∼−0.6 MPa for controls. Berry development was monitored using BRIX degrees.

The studies of Savoi identify the impact of water deficiency on secondary metabolism and genetic regulation in grape berries. In PRJNA348618 by Savoi *et al*. (2017), *Vitis vinifera L.* cv. ‘Merlot’ (clone R3 on SO4 rootstock) was studied over two years, with measurements taken 25 days after anthesis (DAA). Midday stem water potential was kept above −0.6 MPa in controls and maintained between −1.0 and −1.4 MPa in water-stressed plants, with irrigation halted from the fruit set. In PRJNA313234 by Savoi *et al*. (2016) cv. ‘Tocai Friulano’ grafted onto SO4 was studied. Control plants were irrigated to maintain midday stem water potential (ψStem) above −0.8 MPa, while water-stressed plants were irrigated only when ψStem dropped below −1.5 MPa.

The study PRJEB55563, conducted on leaves and roots of cv. ‘Callet’ and cv. ‘Merlot’ over two different years of stress, we utilized only the samples with triplicates. The study classified stress levels based on predawn stem water potential as Mild (-0.6 MPa ≤ Ψstem > -0.9 MPa), Medium (-0.9 MPa ≤ Ψstem > -1.4 MPa), and Extreme (Ψstem ≤ -1.4 MPa) (Rodriguez-Izquierdo *et al*., 2025).

Several other studies were identified in our analysis but were not included in this application for various reasons. PRJNA320028 was discarded due to presenting only two SRA runs, making statistical analysis of the data impossible. PRJNA888237 was not used due to the lack of description in the metadata, making it impossible to identify controls and stressed samples, along with inconsistencies in the amount of data published compared to the analyses presented in the publication (Lin *et al.,* 2013) for leaves and roots of Shine Muscat, Miguang, Red Globe Grape, and Thompson Seedless, related to the number of samples and controls needed for a statistical analysis. PRJNA889459 was discarded because the topic of this study was circRNAs. PRJNA433817 was not used because the study on Pinot Noir was conducted with only an initial RNA-seq control, without adding subsequent control data. For the bioproject PRJNA886074 conducted on Sangiovese, controls were not found within the SRNA metadata. The study PRJNA769649 corresponded to combined copper and water stress. The study PRJNA851974 was discarded because it was conducted under flooded and heatwave stress conditions. The study PRJNA769147 did not describe which samples corresponded to controls and which were stress samples. In the case of PRJNA928668, being a combined experiment of Cabernet Sauvignon and Riesling under heat and drought stress, both together and separately, it was not possible to individualize control samples corresponding to their stress condition according to what was written in the metadata. Lastly, the bioproject PRJNA1206530 was discarded because it involved a methyl jasmonate treatment (grape leaves treated with exogenous MeJa at different times under drought stress).

### Categorization of water stress responses

For the development of a gene expression atlas, the varying levels of water stress tolerance exhibited by different *Vitis* cultivars were taken into account, leveraging the extensive genetic diversity within the *Vitis* genus. This genetic variability influences a wide array of phenotypic traits, including growth, development, stress tolerance, resistance mechanisms, plant architecture, physiology, chemical composition, ecological adaptations, and yield potential (Cadle-Davidson *et al*., 2019). Such diversity results in cultivar-specific responses to water stress, with the *Vitis* genus demonstrating significant variation in traits such as iso/anisohydric behavior (Hochberg *et al*., 2018), embolism resistance (Lamarque *et al.,* 2023), and root structure differences (Bonarota *et al.,* 2024).

However, not all of these factors were incorporated into the analysis. Root architecture was excluded due to insufficient data, and iso/anisohydric classification was not utilized because grapevines exhibit plasticity in their water regulation strategies. Their response may shift between anisohydric and isohydric depending on multiple factors, including the severity of water stress, environmental conditions, cultivar identity, and rootstock influence (Hochberg *et al*., 2018). Instead, the primary criterion for assessing drought tolerance in *Vitis vinifera* cultivars and hybrids was their ability to maintain hydraulic conductivity, a critical factor in plant survival during prolonged or intense drought conditions (Lamarque *et al*., 2023).

To categorize the drought tolerance of different *Vitis* vinifera cultivars, the classification of xylem embolism vulnerability established by Lamarque *et al*. (2023) was applied. According to this classification, the cvs. ‘Chardonnay’ and ‘Callet’ were placed in the medium-to-high vulnerability group, whereas cvs. ‘Cabernet Sauvignon’, ‘Merlot’, and ‘Pinot Noir’ were categorized as having low vulnerability (Lamarque *et al*., 2023). For rootstocks, existing literature provided drought tolerance assessments (Trenti *et al*., 2021; Vitulo *et al.,* 2014; Meggio *et al.,* 2014; Corso *et al.,* 2015), leading to the categorization of *Vitis riparia* as very sensitive, 101-14 as sensitive, and M4 as highly tolerant.

### Search of ABA-related genes and other drought-responsive markers genes

ABA-related genes are many times used as drought marker genes due to their rapid activation or repression in response to drought stress. The grapevine gene catalog (Navarro-Payá *et al.,* 2022) was updated by searching in the literature those identified proteins involved in ABA synthesis and catabolism, its perception and signaling cascades, also including related pathways involved in stomatal closure. A total of 45 key genes related to ABA metabolism and signaling were identified and used to validate the Hydric Stress Atlas tool.

ABA biosynthesis originates from the β-carotene pathway and starts within plastids, where β-carotene is progressively converted into 9′-cis-Neoxanthin and 9′-cis-Violaxanthin. These intermediates are transformed into Xanthoxin by 9′-cis-Epoxycarotenoid Dioxygenase (NCED), a key enzyme in the pathway. Once transported to the cytoplasm, Xanthoxin undergoes oxidation by ABA DEFICIENT (ABA) family enzymes, leading to the formation of Abscisic Acid (ABA). Once synthesized, ABA functions as a phytohormone that regulates plant growth, development, and stress responses. Alternatively, it can be deactivated through catabolism, initiated by hydroxylation at the 8′ position of ABA, a key step mediated by (+)-Abscisic Acid 8′-Hydroxylase (CYP707A), resulting in the formation of the biologically inactive Dihydrophaseic Acid (DPA). Additionally, ABA can be stored in vacuoles via glycosylation, mediated by UGT71C5, and later reactivated through deglycosylation, catalyzed by β-D-Glucopyranosyl Abscisate β-Glucosidase (BG) (Figure 1A; Gong *et al*., 2019; Saito *et al.,* 2004; KEGG; Rossdeutsch *et al.,* 2016).

**Figure 1:**
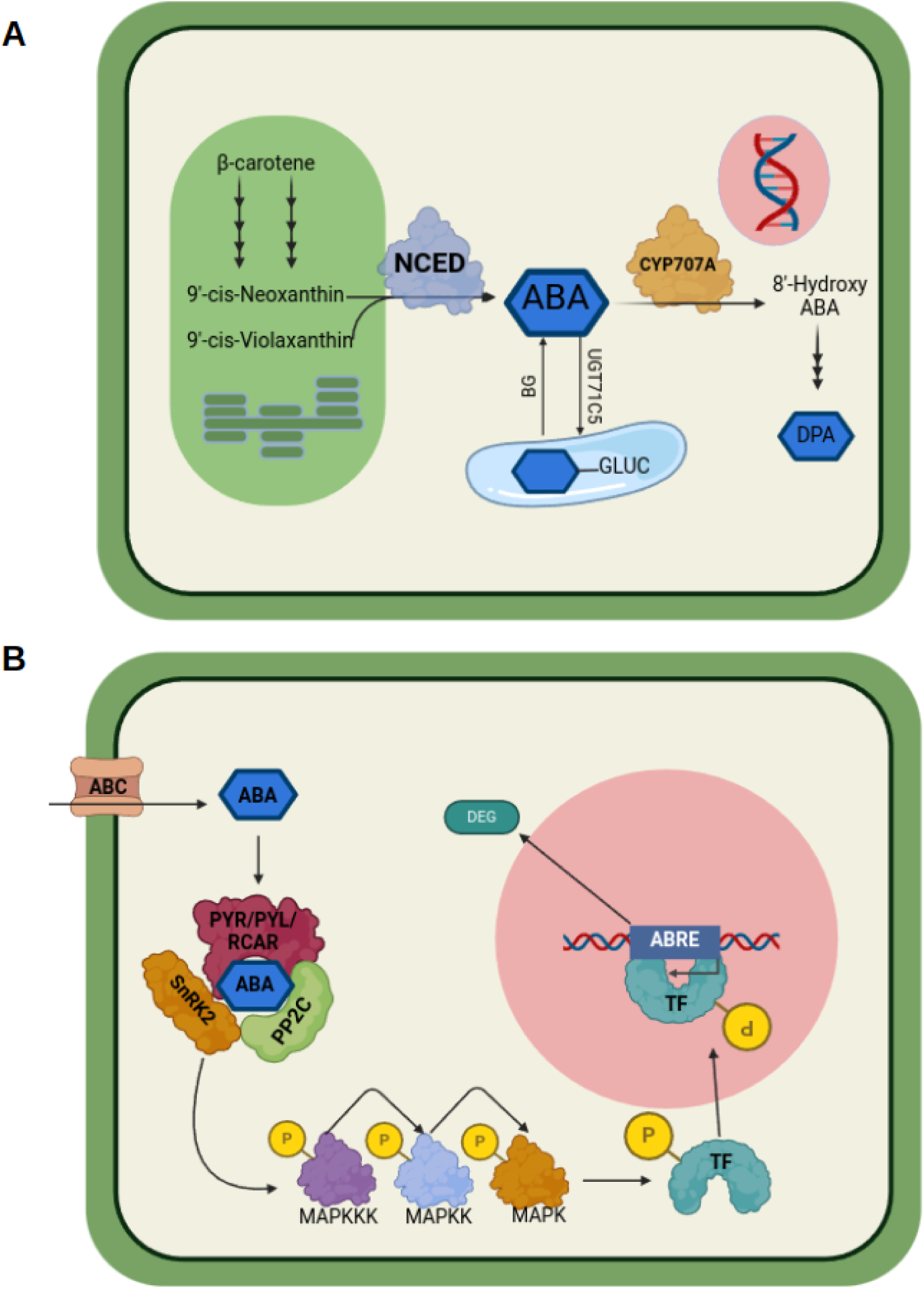
Representation of Abscisic Acid Biosynthesis, Catabolism, and Signaling. (A) A schematic representation of the abscisic acid (ABA) biosynthesis and catabolism pathway. The illustration highlights plastids (green), the nucleus (pink), and the vacuole (blue), all enclosed by the plant cell membrane and wall (green). Key enzymes involved in ABA synthesis (NCED; EC: 1.13.11.51) and catabolism (CYP707A; EC: 1.14.14.137) are emphasized. Additionally, enzymes responsible for ABA storage in vacuoles (UGT71C5/AOG; EC: 2.4.1.263) and its bioavailability (BG; EC: 3.2.1.175) are depicted. (B) A simplified model of the canonical ABA signaling pathway, illustrating the sequence from ABA perception and the MAP kinase cascade to the activation of transcription factors (TFs). These TFs recognize cis-acting ABA-responsive elements (ABREs), leading to the expression of differentially expressed genes (DEGs) in response to ABA. All the proteins named in the figures are described in the supplementary table (A) S1 and (B) S2.

The grapevine genome includes several essential enzymes involved in ABA biosynthesis and degradation. The key enzyme genes for ABA biosynthesis include *VviNCED3*, *VviNCED5*, and *VviNCED6*, which encode 9′-cis-Epoxycarotenoid Dioxygenase (NCED, EC: 1.13.11.51) and catalyze the conversion of 9′-cis-Violaxanthin into Xanthoxin, a crucial precursor in ABA production (Leng *et al*., 2017). The genes coding for these enzymes have been identified on multiple occasions; however, their nomenclature has undergone changes over time (e.g. *VviABCG25* in Cochetel *et al.,* 2020 is *VviABC_G3-1* in Pilati *et al.,* 2017, and He *et al*., 2021). For the release of stored ABA, *VviBG1* plays a central role by hydrolyzing ABA-glucoside in vacuoles, making ABA bioavailable when needed (He *et al.,* 2021). In terms of ABA degradation, the grapevine genome encodes four (+)-Abscisic Acid 8′-Hydroxylase (CYP707A, EC: 1.14.14.137) enzymes—VviHyd1/CYP707A40, VviHyd2/CYP707A41 (Ilc *et al*., 2018), VviHyd3/CYP707A38 (Nicolas *et al*., 2014), and VviHyd4 (Pilati *et al*., 2017; He *et al*., 2021)—which mediate the hydroxylation of ABA, leading to its inactivation as Dihydrophaseic Acid (DPA) (Supplementary Table S1). These enzymes together regulate ABA homeostasis, ensuring a dynamic balance between biosynthesis, storage, and degradation in response to environmental cues.

Once synthesized, the phytohormone ABA regulates a wide range of physiological processes, including plant growth, development, and responses to environmental stress. To elicit these responses, ABA must be perceived and processed within plant cells. This study focuses on two key ABA-mediated pathways: the environmental stress response in guard cells, which leads to stomatal closure, and the canonical ABA signaling pathway, which activates transcription factors that regulate gene expression.

Under water stress conditions, ABA enters cells either through ABC transporters or by passive diffusion in its protonated form (ABAH) (Chen *et al.,* 2019). Inside the cell, ABA is detected by forming the PYR/PYL/RCAR–PP2C–SnRK2 signaling complex, which activates a MAP kinase cascade. This cascade leads to the phosphorylation of bZIP transcription factors, such as AREB/ABF, which recognize ABA-responsive elements (ABRE; PyACGTGGC) in gene promoters. Other transcription factors, including ABI (APETALA2 family) and MYB family members, also contribute to transcriptional regulation under ABA signaling (Figure 1B). In stomatal guard cells, specific proteins such as Open Stomata1 (OST1) play a central role in ABA-mediated responses (Imes *et al*., 2013). However, the study of ABA-dependent transcription factors continues to evolve, revealing additional regulators in this pathway.

In the *Vitis vinifera* PN40024 reference genome, several key genes involved in ABA perception and signaling were identified. Among the ABA transporters, two ABC transporters, *VviABC_G3-1* and *VviABC_G3-2* (Pilati *et al*., 2017; He *et al*., 2021), facilitate ABA uptake. The PYL receptor family, which is responsible for ABA recognition, includes *VviPYL1, VviPYL2, VviPYL4a, VviPYL4b, VviPYL4c, VviPYL8a, VviPYL8b, VviPYL8c*, and *VviPYL11* (Bono *et al*., 2024). Downstream of these receptors, the Protein Phosphatase 2C (PP2C) family, which plays a central role in ABA signal transduction, is represented by *VviPP2C57, VviPP2C58, VviPP2C59, VviPP2C60, VviPP2C61, VviPP2C62, VviPP2C63, VviPP2C64, VviPP2C65, VviPP2C66, VviPP2C67*, and *VviPP2C68* in grapevine (Zhang *et al*., 2021), the PP2C forms a complex with SnRK2 proteins, which recognize ABA molecules to initiate signal transduction via MAP kinases. In *Vitis*, these SnRK2 proteins have been identified as *VviSnRK2A*, *VviSnRK2E*, *VviSnRK2F1*, *VviSnRK2F3*, *VviSnRK2F4* and *VviSnRK2H* (Zhang *et al*., 2021), on the other hand, the MAP kinases identified in *Vitis* include the following *VviMAPKKK14* and *VviMAPKKK15* (Pilati *et al*., 2017; He *et al*., 2021). These components together form the core of the ABA signaling network, regulating gene expression and physiological responses to drought stress in grapevines.

Regarding transcription factors that respond to ABA, the genes *VviABF1* and *VviABF2* (Nicolas *et al*., 2014), *VviBZIP34* (Tu *et al*., 2018), and *VviMYB60* (Galbiati *et al*., 2010) were identified, alongside candidate transcription factors that could be activated in an ABA-dependent manner, such as *VviHB5*, *VviHB7*, *VviMYB143*, and *VviABI5* (Supplementary Table S2). All genes associated with the biosynthesis, catabolism, perception, and signaling pathways were subjected to heatmap analysis for expression changes, utilizing the online tool Hydric Stress Atlas.

### Hydric Stress Atlas app development and validation using marker gene expression

The primary function of the app is to generate heatmaps that illustrate either changes in expression levels (based on log_2_-Fold changes) when compared to control conditions, or single normalized expression per each sample (transcripts per million, TPM). This allows the study of expression patterns across different stress intensities, tissues, and cultivars, with each box representing a comparison to its respective control. Genes are organized in rows, and the app provides an option to cluster them based on the similarity of their expression profiles, facilitating the visualization of expression responses to water stress.

To assess the utility of this expression tool, we examined genes involved in ABA metabolism and signaling. Genes from each functional category were screened for responsiveness in the Hydric Stress Atlas, and those displaying significant changes in expression were selected for visualization. Among them, the genes with most significant downregulation under water stress conditions were *VviPYL1, VviPYL4a, VviPYL4b, VviBG1,* and *VviHyd2*. It was observed that the genes VviBG1 and VviHyd2, involved in the synthesis and catabolism of ABA, exhibited changes in expression depending on the tissue type and genetic background. This expression pattern has been previously reported in drought and heat stress pathways in grapevines by Hewitt et al. (2023). This highlights the critical role of these genes in stress adaptation mechanisms, which may vary based on both environmental and genetic factors. The *VviPYL* genes presented here encode ABA receptors, and their decreased expression upon drought is expected according to Liu and collaborators in 2019. Notably, *VviPYL4a* exhibited expression variation dependent on the genetic background, consistent with findings reported by Rossdeutsch *et al*. (2016) (Figure 2).

**Figure 2.:**
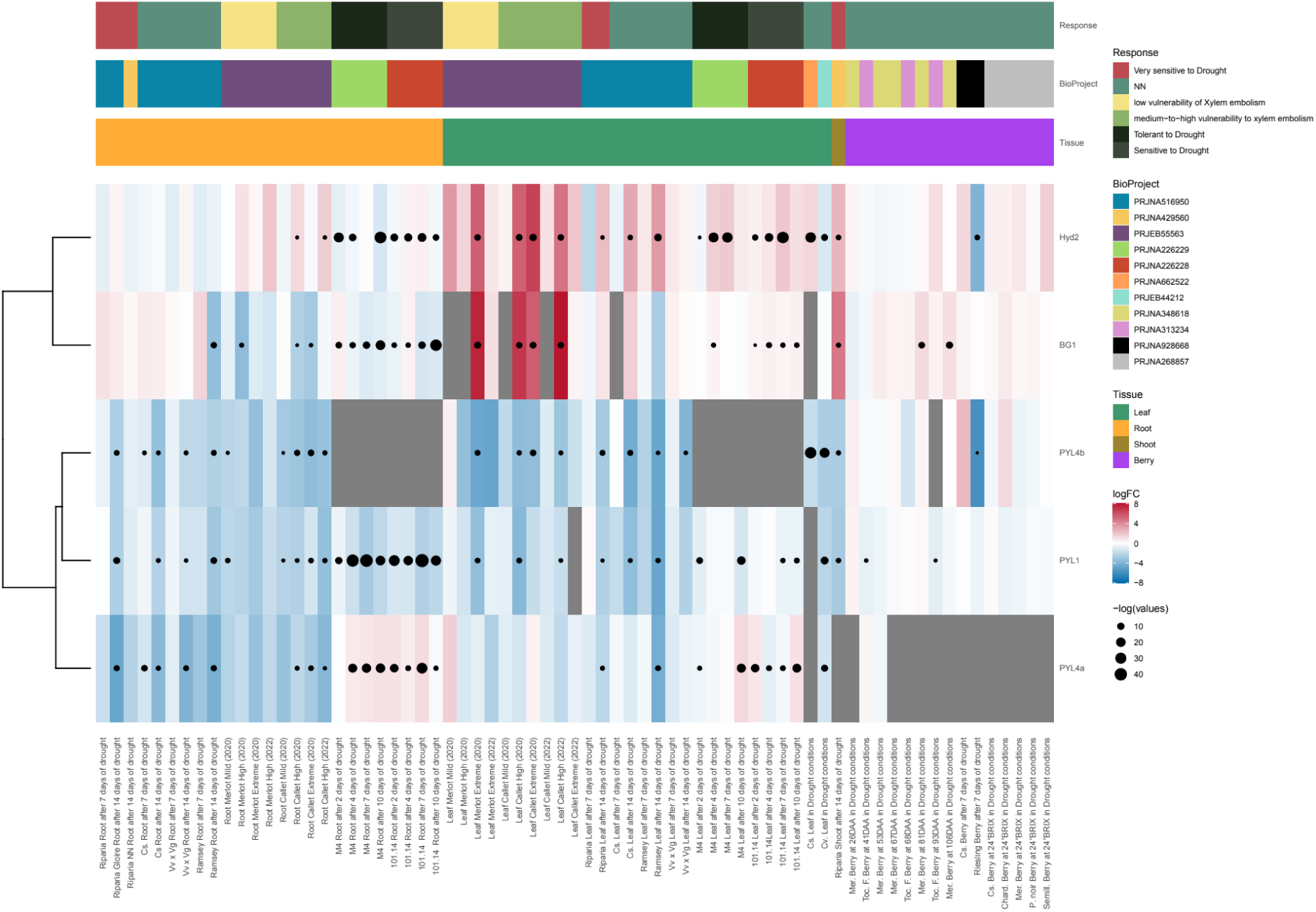
Selection of drought downregulated ABA-related genes. This figure presents a heatmap that visualizes the expression changes of ABA-related genes in response to water stress in grapevine, generated using the Hydric Stress Atlas tool within the PlantaeViz portal. At the top of the figure, three horizontal bars provide contextual metadata information. The first bar categorizes each cultivar or rootstock according to its tolerance to stress, distinguishing between more tolerant and more sensitive varieties. The second bar indicates the source of the data by identifying the corresponding bioproject. The third bar represents the different plant tissues analyzed, such as roots, shoots, leaves, or berries. The main section of the figure consists of the heatmap itself, where each column corresponds to a specific experiment, detailing the tissue, treatment, duration, plant variety, and bioproject involved. Each row represents a specific ABA-related gene. The heatmap uses a color scale to indicate changes in gene expression, with red representing upregulation, blue indicating downregulation genes, and white for no fold change. Gray marks samples where the gene expression is not detected. Boxes may contain dots that denote statistically significant changes, with the dot size proportional to the level of significance. This visualization allows researchers to easily identify expression patterns of key genes under drought conditions across different grape varieties and experimental conditions.

This analysis identified several ABA-related genes that exhibited significant upregulation in response to water stress. Among them, *VviNCED3*, *VviPP2C59*, *VviPP2C60*, *VviPP2C66*, *VviABC_G3-1*, *VviSnRK2F1*, *VviSnRK2F4*, and *VviABF2* showed the most pronounced differences in expression levels. These genes are associated with various key processes within the ABA pathway, including ABA biosynthesis (*NCED3*), as well as the components of the ABA recognition complex, such as the protein phosphatases PP2C and the kinases SnRK2. Additionally, VviABC_G3-1 was identified as an ABA transporter, and VviABF2 as a transcription factor involved in ABA-mediated gene regulation (Nicolas *et al.,* 2014). The analysis also revealed that the expression levels of these genes were notably low in fruit tissues. Furthermore, distinct genetic background-dependent variations were observed for *VviNCED3*, *VviABC_G3-1*, and *VviPP2C66*, indicating that their expression responses to water stress differ depending on the grapevine genotype. These findings highlight key regulatory components of the ABA response under drought conditions and their tissue-specific and genotype-dependent expression patterns (Figure 3).

**Figure 3.:**
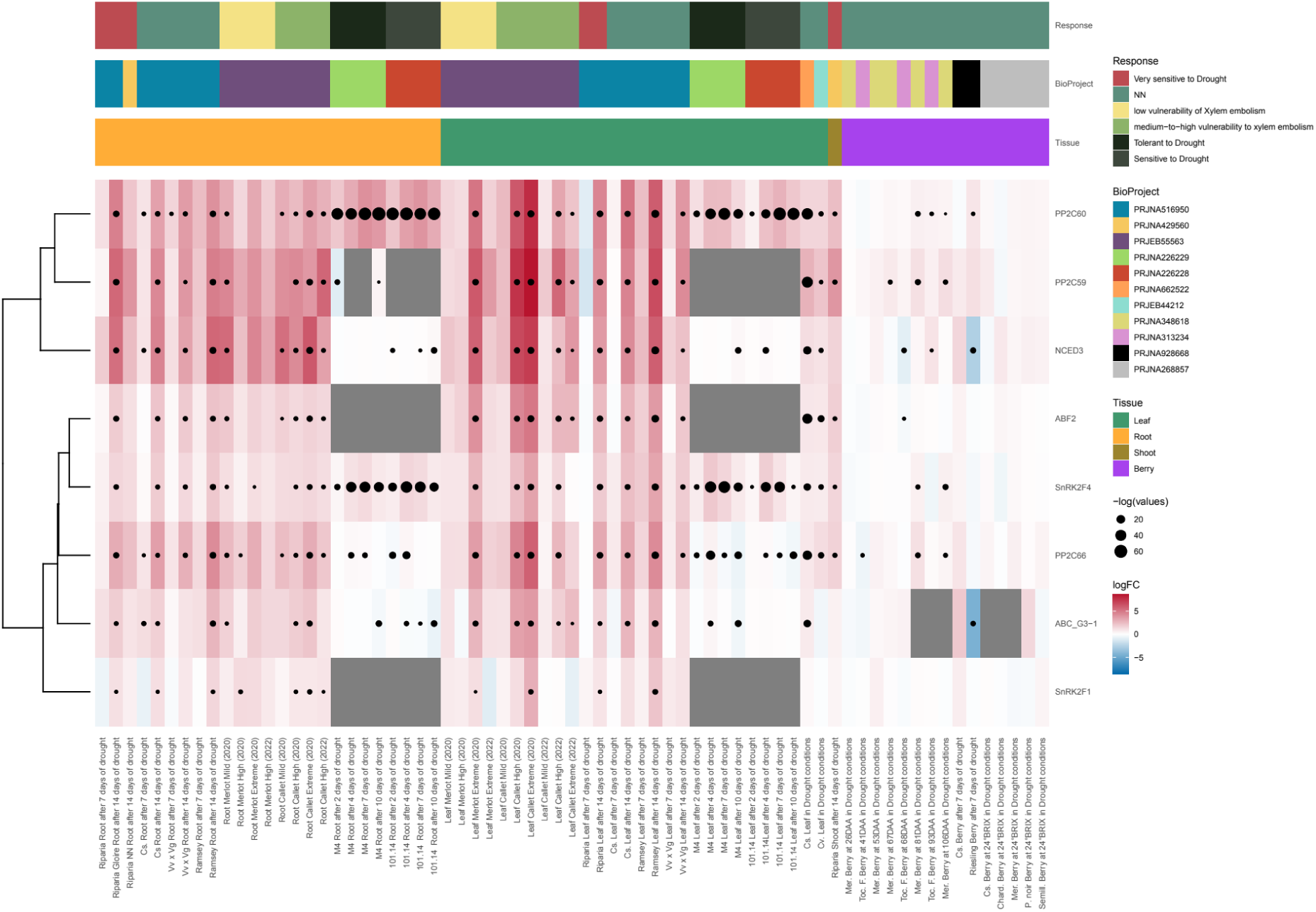
Selection of drought up regulated ABA-related genes. This figure presents a heatmap that visualizes the expression changes of ABA-related genes in response to water stress in grapevine, generated using the Hydric Stress Atlas tool within the PlantaeViz portal. At the top of the figure, three horizontal bars provide contextual metadata information. The first bar categorizes each cultivar or rootstock according to its tolerance to stress, distinguishing between more tolerant and more sensitive varieties. The second bar indicates the source of the data by identifying the corresponding bioproject. The third bar represents the different plant tissues analyzed, such as roots, shoots, leaves, or berries. The main section of the figure consists of the heatmap itself, where each column corresponds to a specific experiment, detailing the tissue, treatment, duration, plant variety, and bioproject involved. Each row represents a specific ABA-related gene. The heatmap uses a color scale to indicate changes in gene expression, with red representing upregulation, blue indicating downregulation genes, and white for no fold change. Gray marks samples where the gene expression is not detected. Boxes may contain dots that denote statistically significant changes, with the dot size proportional to the level of significance. This visualization allows researchers to easily identify expression patterns of key genes under drought conditions across different grape varieties and experimental conditions.

### Exploration of genes related to stomatal development

Another tool found on the Atlas App enables generating box plots displaying log2(FC+1) values for each experiment. Each individual SRA run is represented as a point on the graph, allowing for a detailed examination of drought effects on specific samples. This tool effectively positions control samples alongside stressed samples, making it particularly useful for genes with tissue-specific expression, such as those involved in stomatal development in leaves.

To validate this tool, genes from the stomatal signaling pathway were analyzed, given that stomatal development can be suppressed in response to water stress (Nerva *et al.,* 2023). The objective was to track expression changes in key response genes within green tissues. Stomatal development follows a sequence of fate transitions regulated by specific transcription factors: SPCH controls the transition from protodermal cells to meristemoid cells, MUTE regulates the shift from meristemoid to guard mother cell, and FAMA directs the final differentiation into guard cells (Le *et al.,* 2014; Lau and Bergmann, 2012).

This study specifically examined the transition from protodermal cell to meristemoid cells to identify key genes involved in this developmental response. The transcription factor Hy5 is known to activate the expression of Stomagen, a peptide that inhibits the ERf-TMM receptor, which is responsible for triggering a MAP kinase cascade that ultimately represses the transcription factor SPEECHLESS (SPCH), an essential activator of stomatal development. Consequently, the presence of Stomagen promotes stomatal development. However, other peptides known as EPFs have been identified that activate the ERf-TMM receptor, leading to the inhibition of stomatal development (Wang *et al.,* 2021; Clemens *et al*., 2022; Hunt *et al*., 2010; Figure 4). All the corresponding gene sequences were identified in the *Vitis* genome, confirming the conservation of this regulatory network (Supplementary Table S3).

**Figure 4:**
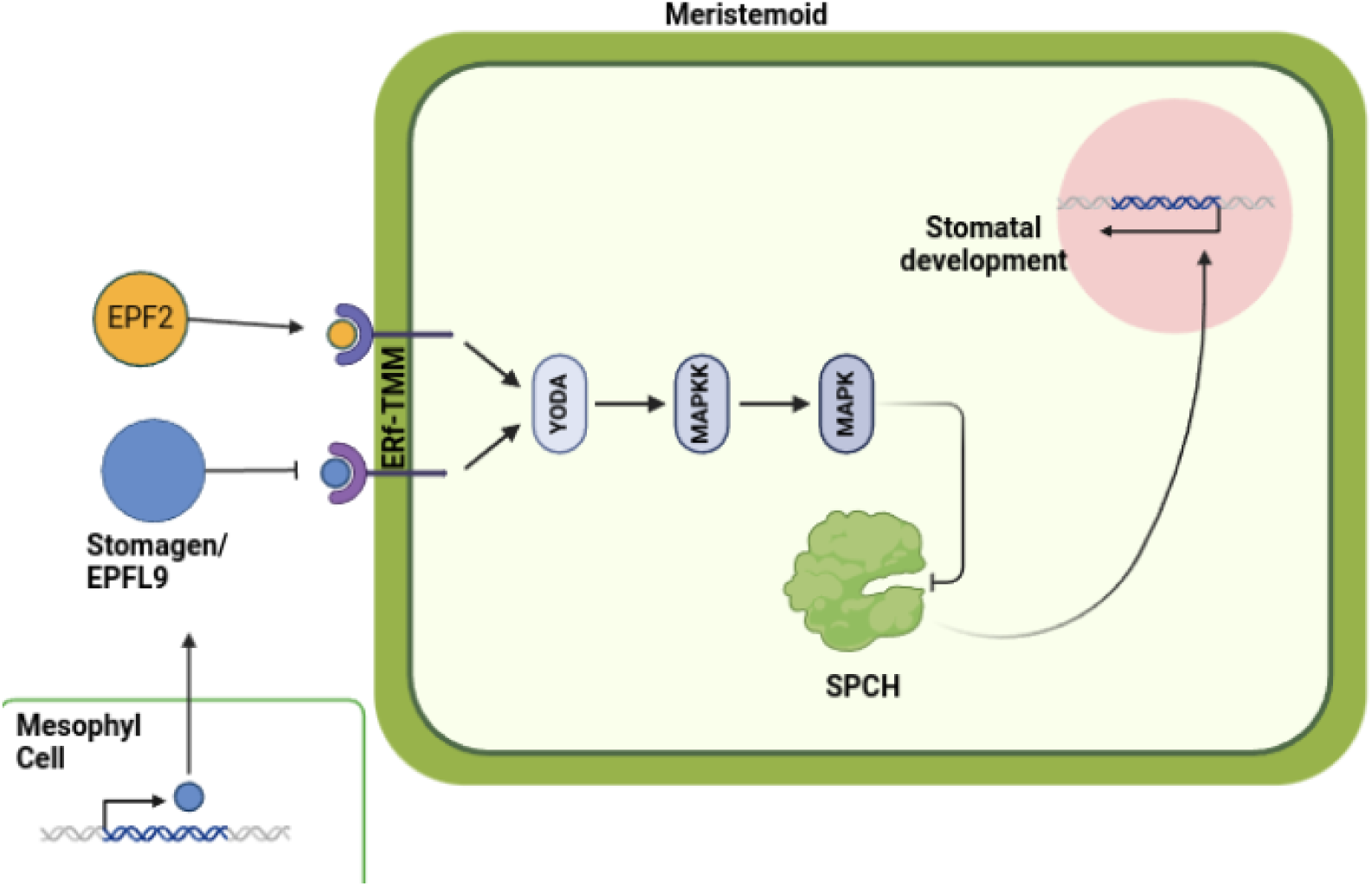
Stomatal development pathway, from protodermal cell to meristemoid. The diagram illustrates the regulatory pathway governing the transition from protodermal cells to meristemoids in stomatal development. It features two green rectangles representing plant cells: one from the mesophyll layer and one from the epidermal layer, emphasizing their chemical communication via the STOMAGEN/EPFL9 peptide. Within the epidermal cell, the ERf-TMM receptor is depicted as the key recognition component of the pathway. Upon ligand binding, the receptor initiates a MAP kinase cascade, leading to the regulation of SPCH (SPEECHLESS), a transcription factor essential for initiating stomatal development. The figure uses arrows with pointed tips to indicate activation events, while blunt arrows represent inhibitory interactions, demonstrating the complex balance between positive and negative regulators in the pathway.

Using the genes described in supplementary Table S3, an expression analysis was conducted using box plots to examine specific expression changes under stress conditions and across different tissues. The analysis provided insights into the regulation of key genes involved in stomatal development and response to environmental cues. For the transcription factor FAMA, which is essential for the final differentiation of guard cells, results showed that its expression is predominantly localized in leaves and shoots. Under mild water stress conditions (between two and four days), a noticeable decrease in FAMA expression was observed. However, as the stress period extended, expression levels returned to match those of the control, suggesting a transient regulatory mechanism that allows the plant to recover from initial stress effects (Figure 5A).

**Figure 5:**
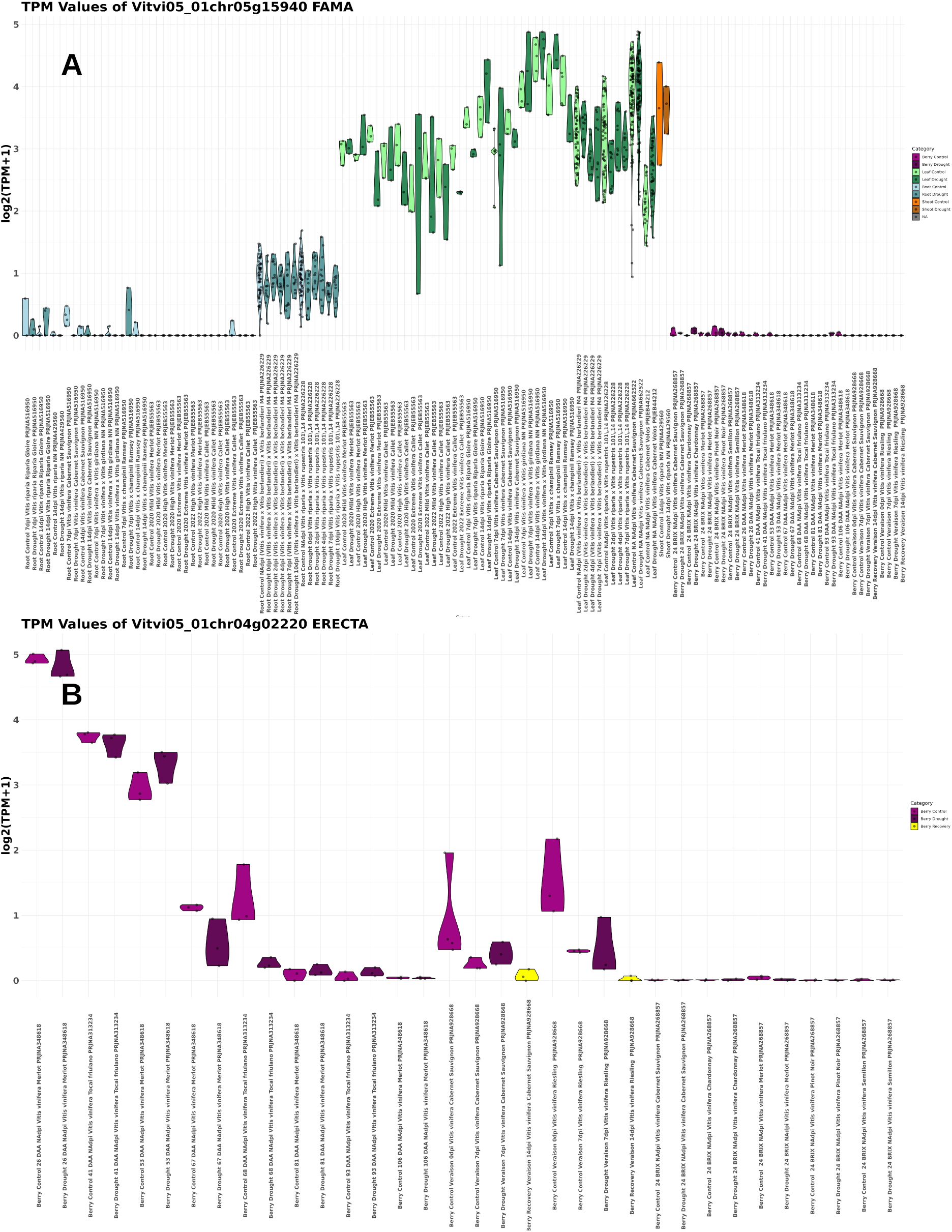
V*viFAMA* and *VviERECTA* expression from the Hydric Stress Atlas. Expression analysis of **(A)** *VviFAMA* (Vitvi05_01chr05g15940) and **(B)** *VviERECTA (*Vitvi05_01chr04g02220) across various tissues in the Hydric Stress Atlas, represented in log2(TPM+1). **(A)** Root samples are shown in blue, leaf samples in green, shoot samples in amber, and berry samples in purple. In all tissues, lighter shades indicate control conditions, while darker shades represent water stress treatments. The X-axis follows the structure: *Tissue, Days of Stress, Organism, Cultivar, Bioproject*, providing an organized overview of expression variations under stress conditions. **(B)** Fruit tissues. The log2(TPM+1) values are displayed with the same color scheme, where lighter shades indicate control conditions, darker shades denote water stress treatments, and yellow indicates recovery condition. The X-axis follows the same structure as in **(A)**, organized from younger to older fruit tissues, progressing from Days After Anthesis (DAA), Veraison, and BRIX°, allowing for a clear visualization of developmental trends in expression.

Regarding the ERECTA receptor, its expression was analyzed in fruit tissues in relation to days after anthesis (DAA). The results revealed that ERECTA expression peaks at 26 DAA, followed by a progressive decline at 41 and 53 DAA, eventually stabilizing at constant levels between 67 and 106 DAA. Beyond this stage, no detectable expression was found at higher maturity levels (measured by BRIX degrees). This trend was consistently observed across various *Vitis vinifera* cultivars included in the Hydric Stress Atlas dataset, highlighting a potential role for ERECTA in early fruit development rather than in later ripening stages (Figure 5B).

### Generation of hydric stress-specific co-expression networks

A drought condition-dependent gene co-expression network (GCNs) was constructed using the datasets detailed in Table 1, incorporating RNA-seq experiments from leaves, shoots, fruits and roots, i.e., tissue-independent (TI). This undirected network consists of 47,636 nodes (genes) and 15,900,784 unique edges (co-expression relationships) and follows a scale-free topology. Performance analysis via EGAD using the v5.1 mapman annotation showed an average AUROC value of 0.69. These hydric stress network can be accessed and downloaded from the GCNs app in the PlantaeViz portal (*VitViz* module; https://plantaeviz.tomsbiolab.com/vitviz/networks/non_agg_gcns/T2T/hydric_stress_TI/), where is possible to generate real-time ‘on the fly’ ontology analyses using the gene-centered networks tab. In addition, an interactive co-expression network can be generated to visualize positive gene interactions in a simplified 3D subnetwork representation, enabling the study of gene interactions under water stress conditions.

**Table 1:**
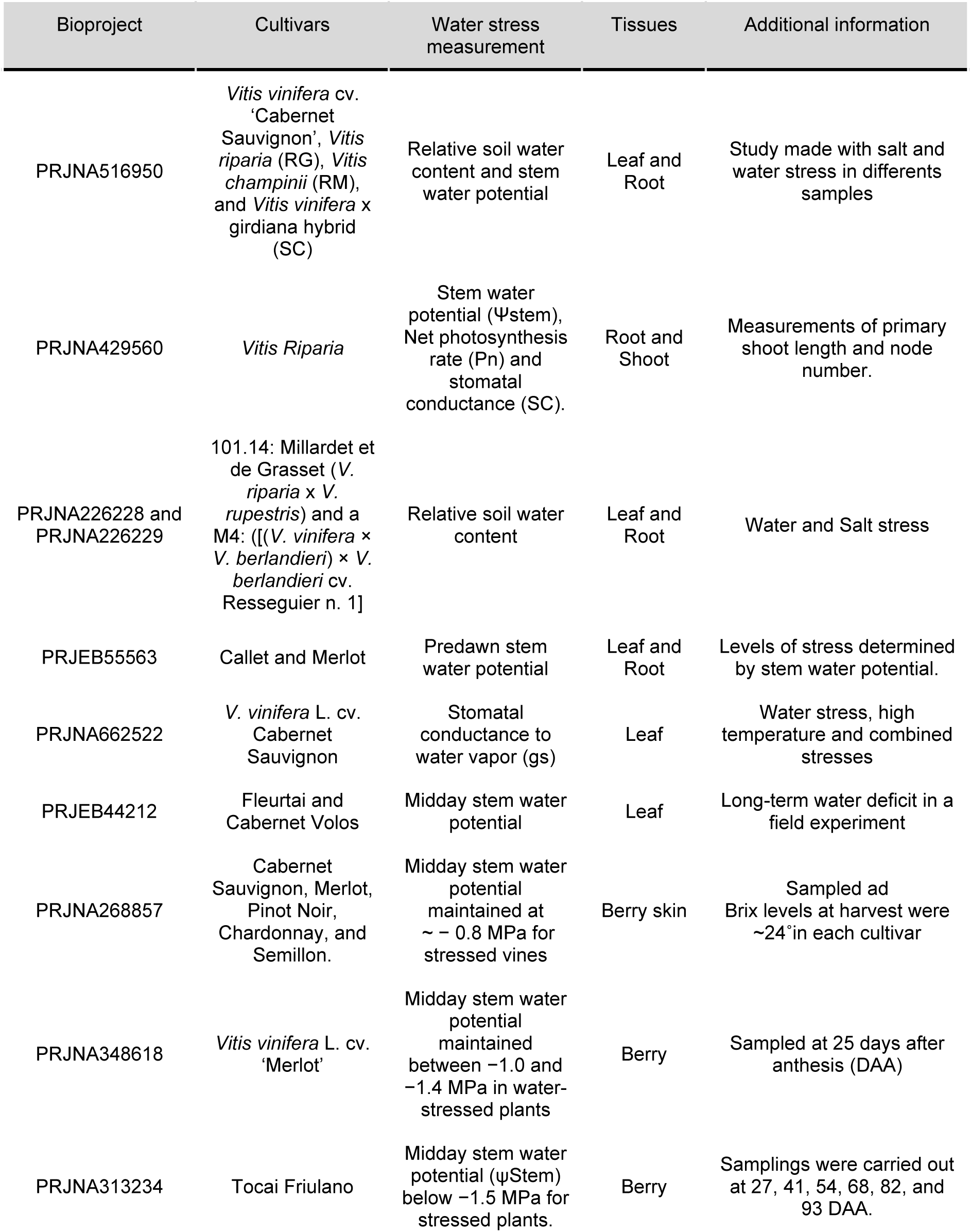
Summary of bioprojects used in the development of the hydric stress atlas. The table below summarizes key details for each project, including the cultivars studied, method employed to quantify and categorize hydric stress, analyzed tissues, and other relevant study information.

#### Case Study 1: Directed analysis on genes with known roles in other species

The drought-responsive GCN was tested to identify transcription factors potentially involved in stomatal development in grapevine, particularly those linked to leaf and guard cell organogenesis. A subset of transcription factors positively associated with stomatal development during drought was compiled (Supplementary Table S4). These transcription factors were analyzed for their co-expression under drought conditions and through category enrichment analyses of their co-expressed genes (top1% genes), searching associations with organogenesis and stress-related processes.

Several transcription factor families previously linked to water stress response or tolerance were identified in the network. Examples include: the lateral organ boundaries domain (LBD) family, a plant-specific TF family involved in grapevine development, berry ripening and stress responses (Grimplet *et al*., 2017), and the GRAS family, playing multifunctional roles in plant growth, development, and resistance to various biotic and abiotic stresses (Waseem *et al*., 2022). The AP2/ERF (APETALA2/ethylene-responsive element binding factors) family was also present. These are key regulators of plant morphogenesis, stress responses, hormone signaling and metabolite regulation (Feng *et al*., 2020). Additionally, the Early Flowering (ELF) TF family, crucial in circadian clock-regulated processes, which integrates environmental light and temperature signals to regulate downstream physiological processes (Zhao *et al*., 2021). Some MYB transcription factors are linked to abiotic stress responses, including drought and cold stress (Fang et al., 2024), as well as abscisic acid (ABA) modulation during water stress (Seo *et al.,* 2009; Abe *et al.,* 2003). Finally, the WRKY TF family is directly involved in drought tolerance, modulating stomatal movement, root architecture, tissue structure, phytohormones and drought-associated metabolic pathways (Li *et al*., 2025). *VviWRKY20* (Vitvi05_01chr07g07840) has been associated with biotic and abiotic tolerance in grapevine (Ma and Yang 2018) andits soybean ortholog has been linked to ABA modulation and drought tolerance (Luo *et al*., 2013), with similar findings in *Amorpha fruticosa* (Li *et al*., 2023) and *Glycine soja* (Tang *et al*., 2014). However, in grapevine, its co-expression network did not indicate direct involvement in ABA modulation.

By contrast, *VviMYB91B*, previously linked to fruit tissue development during early ripening (Ma and Yang, 2018), is overexpressed in early developmental stages, such as buds and seedlings (Fasoli *et al.,* 2012). A gene-centered GCN for *VviMYB91B* revealed its most co-expressed genes were enriched in water stress response-related processes, including: solute transport, protein homeostasis, and carbohydrate metabolism (linked to drought response); signaling cascades (transferases and ubiquitin-proteasome system), and cellular development processes (cell cycle and chromatin organization).

Ontology analysis of the transcription factors listed in Supplementary Table S4 highlighted tissue-specific differences in organogenesis. For example, *VviMYB50* and *VviWRKY44* were exclusively associated with root organogenesis, while *VviLBDIi2* was specific to floral tissues. In contrast, *VviMYB91B* and *VviELF3B* were linked to leaf organogenesis (Figure 6).

**Figure 6:**
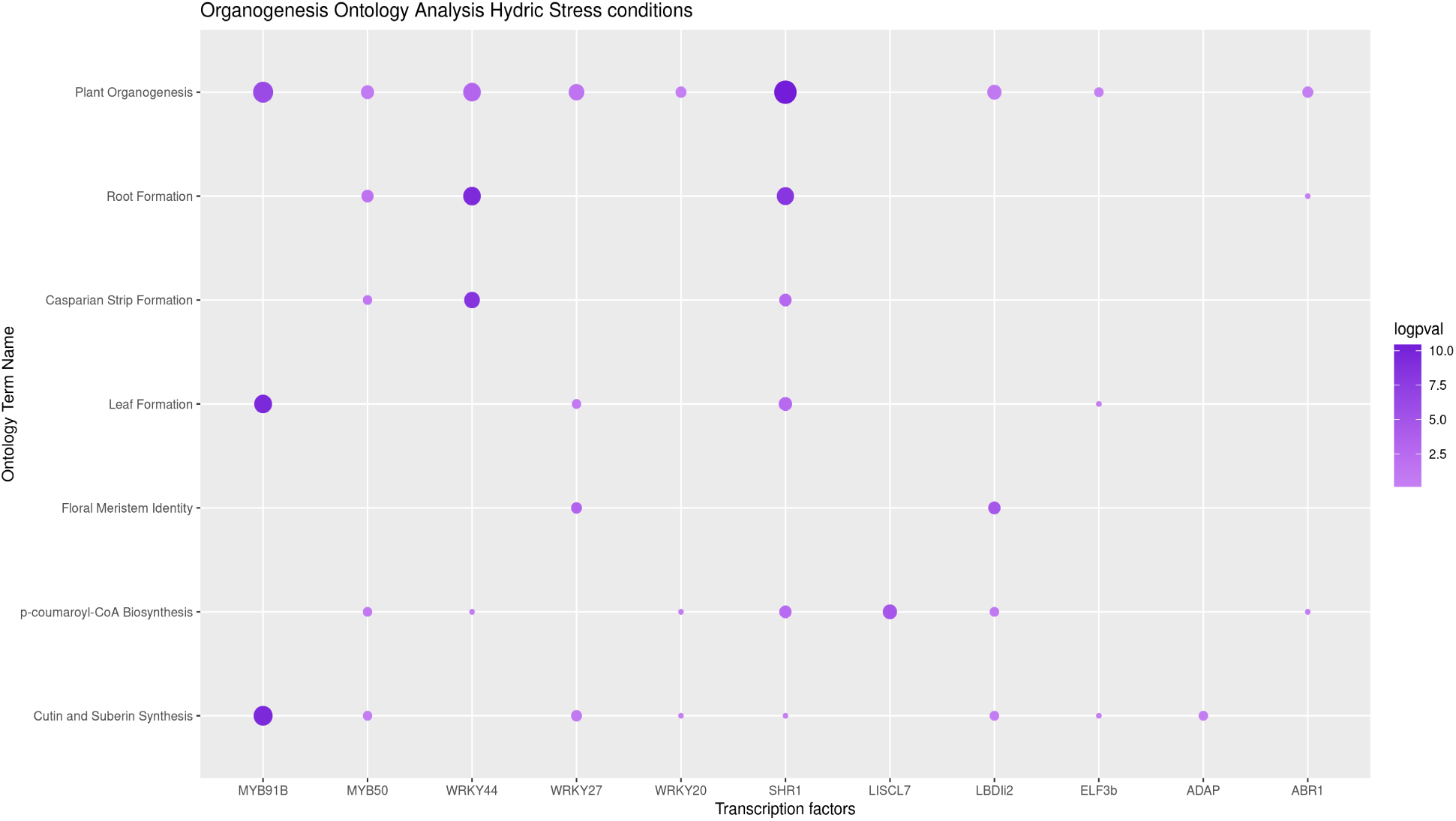
Top Enriched Terms in Transcription factors GCN filtered by Organogenesis processes. The figure displays the enriched gene ontology (GO) terms in the co-expression networks of the transcription factors *VviMYB91B, VviMYB50, VviWRKY44, VviWRKY27, VviWRKY20, VviSHR1, VviLISCL7, VviLBDli2, VviELF3b, VviADAP* and *VviABR1* transcription factors under water stress conditions. The Y-axis represents the different GO terms, while the X-axis shows the log(p-value) of each GO term, corresponding to the number of genes associated with that term in each transcription factor’s co-expression network.

As part of this case study, all transcription factor co-expression networks were examined, generating an Upset Plot to highlight gene intersections across networks and curated gene lists related to ABA biosynthesis (Supplementary Table S1), ABA signaling (Supplementary Table S2), and stomatal development (Supplementary Table S3). The strongest correlation was observed between stomatal development genes and *VviMYB91B*, with three direct associations. Additional correlations were identified, including *VviWRKY20* with *VviWRKY27* (275 shared genes) and *ABR* with *ADAP1* (242 shared genes), among other overlaps (Supplementary Figure S1).

To further explore the role of *VviMYB91B* in a potential stomatal development network, hereafter named *VviMYB91B*-STOMA, an interactive graph was generated, showcasing the top 50 genes most strongly co-expressed with *VviMYB91B* under water stress conditions (Supplementary Table S5). Among these, several key genes related to guard cell development were identified, such as *VviTMM* (Vitvi05_01chr09g08660) and *VviStomagen2* (Vitvi05_01chr18g20880), also known as the epidermal patterning factor. Genes involved in cellular development include Vitvi05_01chr14g25290, which promotes GTP-dependent aminoacyl-tRNA binding during protein biosynthesis, as well as the protodermal factors Vitvi05_01chr12g00520 and Vitvi05_01chr12g07740, which function as epidermal patterning factors. Additionally, two transcription factors from the Squamosa promoter-binding protein-like (SPL) family, *VviSBP3* and *VviSBP14*, were identified and both are associated with plant growth and development (Chen *et al*., 2010). Furthermore, the aquaporin *VviTIP1*-1, known for its role in water deprivation response, was also among the co-expressed genes.

These findings align closely with the ontology enrichment results, further reinforcing the relationships between specific transcription factors and their roles in tissue-specific organogenesis. This GCN-based analysis, guided by the “guilty by association” principle, suggests a central role for *VviMYB91B* in stomatal development under drought conditions. Within the *VviMYB91B* co-expression network, key genes involved in stomatal development were identified, including *VviTMM*, *VviStomagen2*, *VviSPCH* (Vitvi05_01chr16g18130), and *VviERECTA* (Vitvi05_01chr04g02220), as presented in Figure 6 and Table 3. Specifically, during guard cell development, these genes are co-expressed during the transition from protodermal cells to meristemoids, a process regulated by key players such as the stomagen receptor and the MAP kinase cascade effector SPCH, a crucial transcription factor in stomatal development. Together with other highly co-expressed genes under drought conditions, these results strongly support the hypothesis that *VviMYB91B* plays a central role in cell development as a response to drought stress.

#### Case Study 2: Untargeted exploration of highly connected transcription factors and potential regulators of drought responses

After constructing the co-expression networks, a topology analysis was performed using functions from both *igraph* and *influential* R packages, with a particular focus on the integrated value of influence (IVI) metric, which evaluates the significance of a gene within the network. As a result, hub genes were identified as potential key regulators in the water stress response network. The top 100 transcription factors were selected and ranked according to their IVI values (Supplementary Table S6). These transcription factors were further examined by constructing a GCN subnetwork using the D3 subnetwork tool, revealing four distinct clusters (Figure 8).

Figure 7 provides a 2D representation of the 3D interactive co-expression networks generated using the networkD3 library in R. The top 12 transcription factors (TFs) with the highest Integrated Value of Influence (IVI) were identified, including *VvibZIP20, VvibZIP23, VviMYB150, VviNAC72, VviARF6, VviZFP3, VviABI4, VviLBDIc9, VviDRNL, VviLAS1, VviMYB160,* and *VviZFP15*. This representation facilitates the study of GCNs and provides insights into the potential regulatory roles of key transcription factors under water stress conditions.

**Figure 7:**
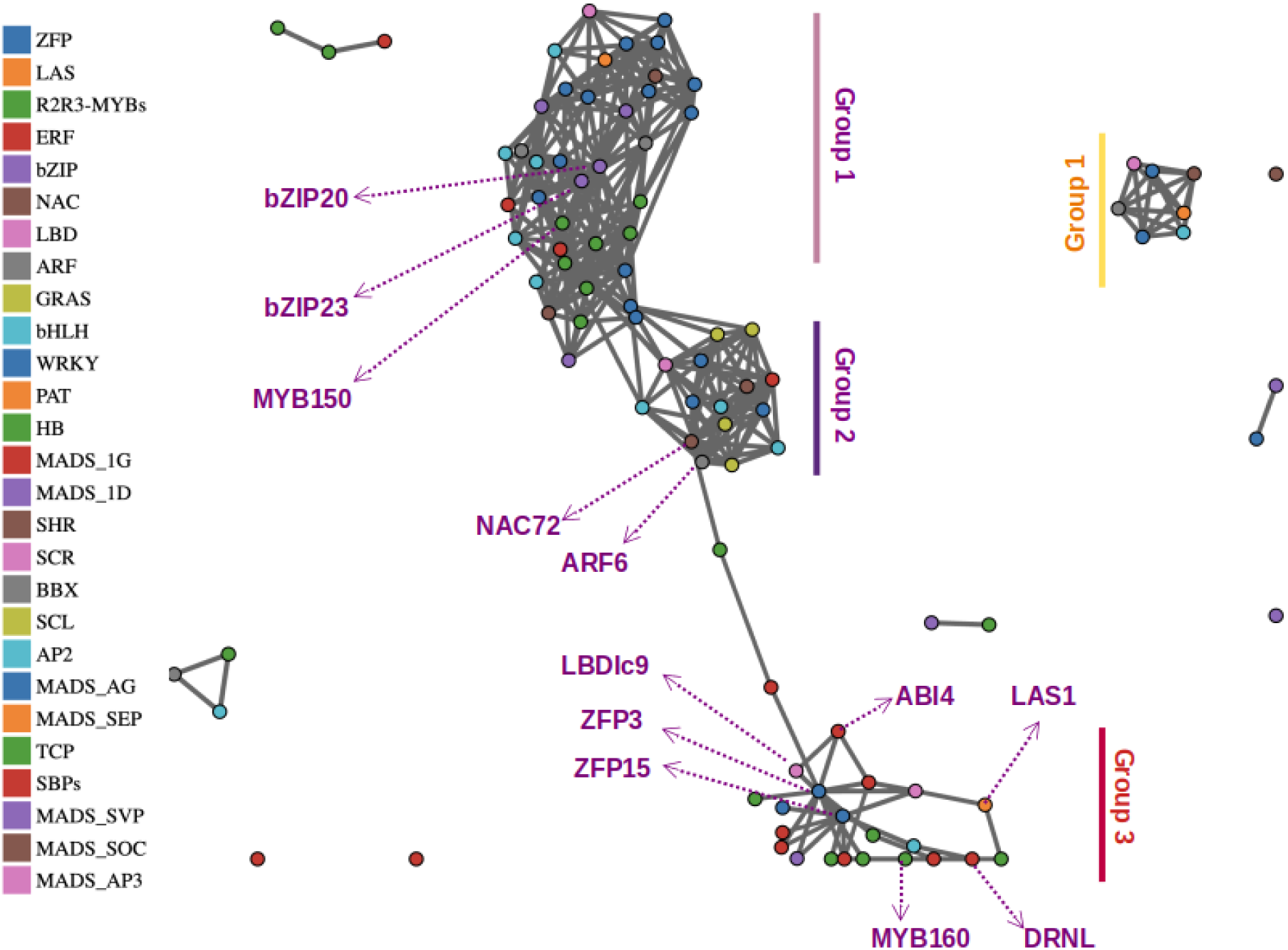
TOP100 TFs by integrated value of influence (IVI). Network D3 visualization of the TOP100 TFs ranked by IVIs using the hydric stress tissue independent (TI) network app within PlantaeViz. A force-directed representation based on how the selected genes co-express with each other is derived from the number and weight of edges between nodes. The legend identifies the transcription factor groups used in the analysis, while the central figure illustrates how all displayed genes are directly connected in the network. In the visualization, four distinct co-expression network groups are highlighted, each labeled from one to four for clarity and organization.

To streamline the analysis, the top 12 TFs from this ranked list were selected for an in-depth study within these clusters, providing deeper insights into their roles and interactions within the co-expression networks. Our analysis demonstrates the functional divergence among the clusters, highlighting their specialized roles in regulating stress responses. Cluster 4 is enriched in functions associated with floral organ development, with key genes such as *VviSEP3*, *VviAP3b*, *VviAP2* (APETALA family), *VviAG1*, and *VviAGL15a* (AGAMOUS family), which are strongly linked to floral structure formation and differentiation. This underscores Cluster 4’s specialized role in floral organogenesis. In contrast, Clusters 1 to 3 were analyzed in detail, focusing on the top 12 TFs ranked by their IVI values. The co-expression network clusterization was consistent with the MapMan ontology analysis, revealing shared ontologies related to solute transport, carbohydrate metabolism, and aquaporin channels, all crucial for plant adaptation to water stress.

Cluster 1 is characterized by ontologies associated with brassinosteroids, jasmonates, root organogenesis, and gravity sensing, all closely linked to root system architecture (RSA) adjustments, a key adaptive mechanism under water stress conditions. Cluster 2 is enriched in abscisic acid (ABA)-related ontologies, reinforcing ABA’s key regulatory role in water stress responses. Cluster 3 exhibits unique redox balance and oxidative stress response ontologies, highlighting critical components for cellular homeostasis and survival under drought conditions. Together, these findings emphasize the importance of these transcription factors and their associated networks in unraveling the complex mechanisms plants use to cope with water stress (Figure 8).

**Figure 8:**
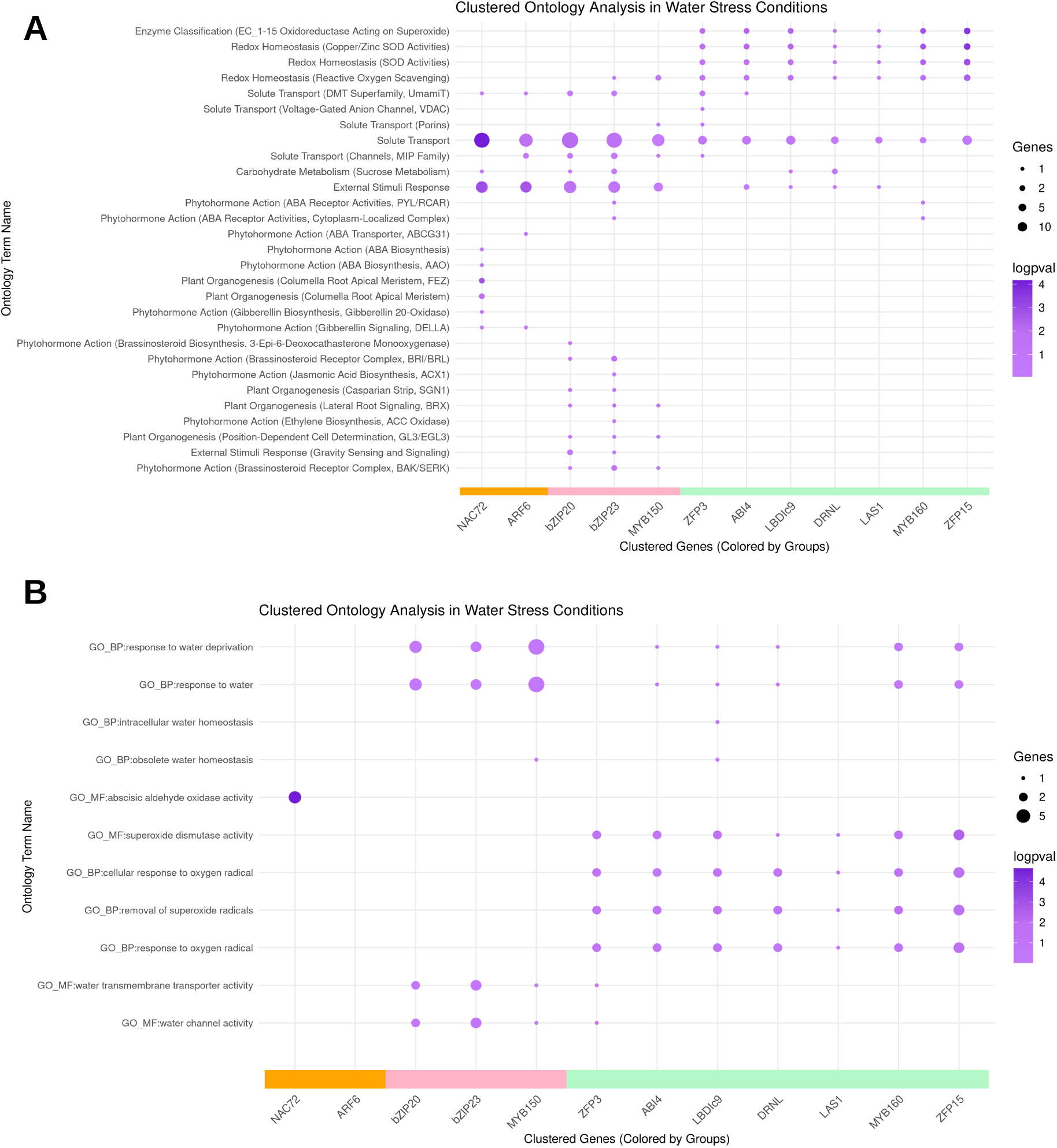
Top (A) MapMan and (B) GO enriched terms in Top 12 transcription factors. This bubble chart provides a comprehensive visualization of the relationship between MapMan ontological terms associated with water stress responses and the top 12 transcription factors (*VvibZIP20, VvibZIP23, VviMYB150, VviNAC72, VviARF6, VviZFP3, VviABI4, VviLBDIc9, VviDRNL, VviLAS1, VviMYB160,* and *VviZFP15*). The Y-axis lists the water stress-related ontological terms, while the X-axis displays the transcription factors, separated into three clusters based on ontological term differences, that match with the GCN clusters. These clusters are color-coded orange for group two, pink for group one and green for group three enhancing visual clarity. At the core of the chart, each bubble represents genes within the respective co-expression networks (GCNs) of the transcription factors associated with a given ontological term. The bubble size correlates with the number of genes within the GCNs, whereas the purple intensity reflects the negative log(p-value) of each gene group, as indicated in the legend. This visualization effectively captures gene regulation patterns linked to water stress responses, emphasizing the key enriched ontological terms within each transcription factor’s network.

## Discussion

Addressing climate change and water stress in grapevines requires innovative breeding strategies to enhance drought resilience. While conventional breeding techniques (CBTs), such as QTL mapping and GWAS, have identified key drought tolerance traits, they involve long breeding cycles. In contrast, new breeding techniques (NBTs), particularly CRISPR-Cas9, offer precise genome editing to accelerate adaptation. Despite challenges like prolonged regeneration times, significant advancements in grapevine genetic editing have been made (Magon *et al.,* 2023; Malnoy *et al.,* 2016). Additionally, systems biology approaches provide powerful tools to dissect grapevine stress responses. Developing F.A.I.R. (Findable, Accessible, Interoperable, and Reusable) transcriptomic resources, such as the Hydric Stress Atlas by PlantaeViz, is crucial for advancing biotechnological research.

This study introduced the PlantaeViz platform, which facilitates transcriptomic analysis of *Vitis* under drought conditions. Gene expression heatmaps revealed key genes in ABA biosynthesis and signaling, with *VviPYL1*, *VviPYL4a*, *VviPYL4b*, *VviBG1*, and *VviHyd2* downregulated, while *VviNCED3*, *VviPP2C59*, *VviPP2C60*, *VviPP2C66*, *VviABC_G3-1*, *VviSnRK2F1*, *VviSnRK2F4*, and *VviABF2* were upregulated. These genes serve as potential hydric stress markers in *Vitis vinifera*, adaptable to different genetic backgrounds.

Two additional tools allowed for deeper transcriptomic analysis using stomatal development as a model. The first, an RNA-seq data explorer, showed how drought affects gene expression patterns, exemplified by *VviFAMA*, a key regulator of guard cell development, and *VviERECTA*, whose expression varies with fruit development. The second tool, a condition-dependent co-expression network, identified TF families highly interactive under drought, including MYBs, WRKYs, GRASs, AP2, LBD, and ELF. Notably, *VviMYB91B* was co-expressed with key stomatal development genes, suggesting a regulatory role in drought-induced cellular development. A topology analysis of the Hydric Stress network identified the most influential transcription factors organized into three major co-expression clusters corresponding to key water stress pathways: ABA signaling, oxidative stress regulation, and root system architecture (RSA). A fourth cluster, linked to floral organ development, highlighted potential water-stress-induced flowering mechanisms mediated by APETALA genes. The transcription factors identified in Supplementary Table S6 represent candidate regulators of water stress responses in grapevine, warranting further functional validation.

This comprehensive transcriptomic toolkit provides valuable insights into gene regulation under hydric stress and a foundation for future studies, including pan-transcriptomics, molecular marker identification, and genetic modifications. Ultimately, such advancements will safeguard the centuries-old tradition of winemaking, now at risk due to climate change.

## Methods

### Transcriptomic Analysis

A comprehensive *in-silico* analysis was conducted on all available RNA-Seq raw data related to drought stress, utilizing a thorough bibliographic collection from NCBI papers and the SRA database. The specific search query employed was: “(grapevine [Title/Abstract] OR grape [Title/Abstract] OR Vitis [Title/Abstract] OR V. vinifera [Title/Abstract]) AND (drought OR water stress OR water deficiency OR water deficit stress) AND (transcriptome [Title/Abstract] OR transcript profiling [Title/Abstract] OR mRNA expression [Title/Abstract] OR RNA-Seq [Title/Abstract] OR RNASeq [Title/Abstract] OR RNA Sequencing [Title/Abstract] OR RNA-Sequencing [Title/Abstract])”. This query resulted in the identification of over 80 papers. Each paper was meticulously reviewed, filtered, and organized by Bioproject for re-analysis using the SRA database. Ultimately, a list of 10 bioprojects comprising 1203 RUN IDs was compiled. The initial data organization was quite disordered. Subsequently, the data was organized to eliminate non triplicated runs and sorted based on treatment, tissue, type of stress, developmental stage, cultivar, organism, and unspecified data, culminating in a final dataset of 1107 RUN IDs for further computational analysis. The selected bioprojects include: PRJNA516950 (Cochetel *et al.,* 2020), PRJNA429560 (Kahdka *et al.,* 2019), PRJNA226228 (Vitulo *et al.,* 2014; Meggio *et al.,* 2014), PRJNA226229 (Corso *et al.,* 2015; Meggio *et al.,* 2014), PRJNA662522 (Tan *et al.,* 2023), PRJEB44212, PRJNA268857 (Ghan *et al.,* 2015), PRJNA348618 (Savoi *et al.,* 2017), and PRJNA313234 (Savoi *et al.,* 2016). The resulting runs were downloaded from SRA using the *prefetch* command with defailt parameters. Fastq files were generated using *fastq-dump* using default parameters. Illumina reads were trimmed with Fastp software (Chen *et al.,* 2018) and aligned to the grapevine 12X.v2 PN40024 genome using STAR (Dobin *et al.,* 2013). Raw counts were generated with FeatureCounts (Liao *et al.,* 2014) using the VCost.v3 genome annotation. Differential expression analysis was conducted using the LIMMA R package (Ritchie *et al.,* 2015), and FPKM normalization was performed with the DESeq2 package (Love *et al.,* 2014). Genes were classified as differentially expressed if the adjusted p-value was below 0.05.

The analysis results were deployed as an app on the VitViz platform, showcasing a heatmap where cell color represents logFC for each comparison, and dot size corresponds to - log(pvalue). Users can upload a file with genes of interest for display, with each experiment described specifically by tissue, water stress intensity, cultivar/rootstock, water stress tolerance level, bioproject, and developmental stage in the case of berries.

### Violin plot analysis

Violin plot analyses were conducted in R using the “tidyverse”, “readODS”, and “patchwork” packages. The TPM results were used to perform the normalization calculation log(TPM+1), which was then reorganized and presented in the form of a violin plot using the “ggplot” package.

### Condition-dependent Gene Co-Expression Network (GCN)

GCN generation commenced by selecting SRA studies based on tissue origin. A total of 476 runs were from ABI solid platform data and 630 from Illumina. Illumina reads were downloaded from SRA, trimmed using fastp (with default parameters + -- cut_front_window_size 1 --cut_front_mean_quality 30 --cut_front cut_tail_window_size 1 -- cut_tail_mean_quality 30 --cut_tail -l 20), and aligned to the PN40024.v5 genome using STAR (with default parameters + --runMode alignReads --limitOutSJcollapsed 8000000 -- limitIObufferSize 220000000), following the pipeline by Orduña *et al*. (2023). Count matrices were obtained using FeatureCounts (-t “exon” -C -g “featurecounts_id”) with the PN40024.v5.1 annotation, summarizing counts at the gene level even when counting reads within exons. ABI solid platform reads followed the same pipeline except using Cushaw3 as the aligner with default parameters. All SRA runs used in a particular GCN were combined into a single count matrix, and runs with less than 4 million successfully aligned reads were excluded. The overall method was executed as previously described for single or non-aggregated approaches (Santiago *et al.,* 2024; Orduña *et al*., 2023). Raw count matrices from the SRA studies were normalized to TPM, discarding genes with less than 0.5 TPMs in every input run. Pearson’s correlation coefficients for each gene against all other remaining genes were calculated and ranked in descending order. These ranked Pearson’s correlation coefficient (PCC) values were used to compute the highest reciprocal ranking (HRR) matrix (Mutwil *et al*., 2010), considering only the top 479 ranked genes (approximately 1% of all PN40024.v5.1 gene models), using the formula: HRR(A,B) = max(rank(A,B), rank(B,A)). Finally, as a noise-filtering step, only the top 479 values for each gene were included in the final network construction. The visualization of the co-expression network was made by using the networkD3 package in R. Network performance was assessed using the EGAD R package, where an average area under receiver operating characteristic curve (AUROC) was calculated for 10 repetitions of the run_GBA function.

## Acknowledgements

This work was supported by the Edmund Mach Foundation in collaboration with the University of Trento, the Institute for Integrative System Biology at the University of Valencia, and the Research and Innovation Center of Concha y Toro Vineyards. This work was also supported by grant PID2021-128865NB-I00 awarded to J.T.M from the Ministerio de Ciencia, Innovación y Universidades (MCIU, Spain), Agencia Estatal de Investigación (AEI, Spain), and Fondo Europeo de Desarrollo Regional (FEDER, European Union). A.S. was supported by the PROMETEO scholarship (PROMETEO/2021/056-01) from the Generalitat Valenciana (GVA). A.V.V. was also supported by the Cost Action CA17111 “Integrape” through the grant E-COST-GRANT-CA17111-2f2cdd5e titled “In-silico analyses using the 40X-improved *Vitis vinifera* genome fourth assembly for identifying hub genes,” which was used for the generation of the Hydric Stress Atlas. The generation of the networks and all the bioinformatic analyses were performed on the HPC cluster Garnatxa at the Institute for Integrative Systems Biology (I2SysBio, UV-CSIC, Spain). We also thank Alberto Rodríguez and David Carrasco (CBGP, Madrid) for data sharing.

## Author contributions

J.T.M., M.M., F.G and A.V.V. conceived the work. A.V.V. performed the data search and meta-data curation, and together with J.T.M. designed the case study examples. A.V.V., D.N.P., A.S. and P.S. conducted the transcriptomic analyses, GCN generation and app development within VitViz. A.V.V. and D.N.P. analysed the results. All authors reviewed the manuscript.

## Data availability statement

In compliance with FAIR (findable, accessible, interoperable, and reusable) guidelines, data has been made accessible through the applications themselves, both in raw form and in the various processed subsets that can be obtained from the various online tools. All the data presented in this study are available for use at the VitViz module at the following links https://plantaeviz.tomsbiolab.com/vitviz/hydric_atlas/ for the hydric expression atlas, and https://plantaeviz.tomsbiolab.com/vitviz/networks/non_agg_gcns/T2T/hydric_stress_TI/ for the non aggregated gene co-expression network. Gene catalogs were updated with ABA and stomata-related genes, available at https://plantaeviz.tomsbiolab.com/vitviz/catalogue/.

## Additional Information (including a Competing Interests Statement)

The authors declare that they have no known competing financial interests or personal relationships that could have appeared to influence the work reported in this paper.

## REFERENCES

Abe, H., Urao, T., Ito, T., Seki, M., Shinozaki, K., & Yamaguchi-Shinozaki, K. (2002). Arabidopsis AtMYC2 (bHLH) and AtMYB2 (MYB) Function as Transcriptional Activators in Abscisic Acid Signaling. The Plant Cell, 15(1), 63–78. 10.1105/tpc.006130

Bonarota, M.-S., Toups, H. S., Bristow, S. T., Santos, P., Jackson, L. E., Cramer, G. R., & Barrios-Masias, F. H. (2024, March). Drought response and recovery mechanisms of grapevine rootstocks grafted to a common Vitis vinifera scion. Plant Stress. Elsevier BV. Retrieved from 10.1016/j.stress.2024.100346

Bono, M., Ferrer-Gallego, R., Pou, A., Rivera-Moreno, M., Benavente, J. L., Mayordomo, C., Deis, L., Carbonell-Bejerano, P., Pizzio, G. A., Navarro-Payá, D., Matus, J. T., Martinez-Zapater, J. M., Albert, A., Intrigliolo, D. S., & Rodriguez, P. L. (2024). Chemical activation of ABA signaling in grapevine through the iSB09 and AMF4 ABA receptor agonists enhances water use efficiency. Physiologia Plantarum, 176(6). 10.1111/ppl.14635

Cadle-Davidson, L., Londo, J., Martinez, D., Sapkota, S., & Gutierrez, B. (2019). From Phenotyping to Phenomics: Present and Future Approaches in Grape Trait Analysis to Inform Grape Gene Function. Compendium of Plant Genomes. Springer International Publishing. Retrieved from 10.1007/978-3-030-18601-2_10

Cerda, A., & Alvarez, J. M. (2023, November 16). Insights into molecular links and transcription networks integrating drought stress and nitrogen signaling. New Phytologist. Wiley. Retrieved from 10.1111/nph.19403

Clemens, M., Faralli, M., Lagreze, J., Bontempo, L., Piazza, S., Varotto, C., Malnoy, M., Oechel, W., Rizzoli, A., & Costa, L. D. (2022). VvEPFL9-1 Knock-Out via CRISPR/Cas9 Reduces Stomatal Density in Grapevine. Frontiers In Plant Science, 13. 10.3389/fpls.2022.878001

Cochetel, N., Ghan, R., Toups, H. S., Degu, A., Tillett, R. L., Schlauch, K. A., & Cramer, G. R. (2020, February 4). Drought tolerance of the grapevine, Vitis champinii cv. Ramsey, is associated with higher photosynthesis and greater transcriptomic responsiveness of abscisic acid biosynthesis and signaling. BMC Plant Biology. Springer Science and Business Media LLC. Retrieved from 10.1186/s12870-019-2012-7

Corso, M., Vannozzi, A., Maza, E., Vitulo, N., Meggio, F., Pitacco, A., Telatin, A., et al. (2015, June 2). Comprehensive transcript profiling of two grapevine rootstock genotypes contrasting in drought susceptibility links the phenylpropanoid pathway to enhanced tolerance. Journal of Experimental Botany. Oxford University Press (OUP). Retrieved from 10.1093/jxb/erv274

Chen, K., Li, G., Bressan, R. A., Song, C., Zhu, J., & Zhao, Y. (2019). Abscisic acid dynamics, signaling, and functions in plants. Journal Of Integrative Plant Biology, 62(1), 25–54. 10.1111/jipb.12899

Chen, X., Zhang, Z., Liu, D., Zhang, K., Li, A., & Mao, L. (2010). SQUAMOSA Promoter-Binding Protein-Like Transcription Factors: Star Players for Plant Growth and Development. Journal Of Integrative Plant Biology, 52(11), 946–951. 10.1111/j.1744-7909.2010.00987.x

Djennane, S., Gersch, S., Le-Bohec, F., Piron, M.-C., Baltenweck, R., Lemaire, O., Merdinoglu, D., et al. (2023, December 9). CRISPR/Cas9 editing of Downy mildew resistant 6 (DMR6-1) in grapevine leads to reduced susceptibility to Plasmopara viticola. (M. Höfte, Ed.)Journal of Experimental Botany. Oxford University Press (OUP). Retrieved from 10.1093/jxb/erad487

Dobin, A., Davis, C. A., Schlesinger, F., Drenkow, J., Zaleski, C., Jha, S., Batut, P., et al. (2012, October 25). STAR: ultrafast universal RNA-seq aligner. Bioinformatics. Oxford University Press (OUP). Retrieved from 10.1093/bioinformatics/bts635

Dong, Y., Duan, S., Xia, Q., Liang, Z., Dong, X., Margaryan, K., Musayev, M., et al. (2023, March 3). Dual domestications and origin of traits in grapevine evolution. Science. American Association for the Advancement of Science (AAAS). Retrieved from 10.1126/science.add8655

Fang, L., Wang, Z., Su, L., Gong, L., & Xin, H. (2023). Vitis Myb14 confer cold and drought tolerance by activating lipid transfer protein genes expression and reactive oxygen species scavenge. Gene, 890, 147792. 10.1016/j.gene.2023.147792

Fasoli, M., Dal Santo, S., Zenoni, S., Tornielli, G. B., Farina, L., Zamboni, A., Porceddu, A., et al. (2012, September 1). The Grapevine Expression Atlas Reveals a Deep Transcriptome Shift Driving the Entire Plant into a Maturation Program. The Plant Cell. Oxford University Press (OUP). Retrieved from 10.1105/tpc.112.100230

Feechan, A., Jermakow, A. M., Torregrosa, L., Panstruga, R., & Dry, I. B. (2008). Identification of grapevine MLO gene candidates involved in susceptibility to powdery mildew. Functional Plant Biology. CSIRO Publishing. Retrieved from 10.1071/FP08173

Feng, K., Hou, X., Xing, G., Liu, J., Duan, A., Xu, Z., Li, M., Zhuang, J., & Xiong, A. (2020). Advances in AP2/ERF super-family transcription factors in plant. Critical Reviews In Biotechnology, 40(6), 750–776. 10.1080/07388551.2020.1768509

Galbiati, M., Matus, J. T., Francia, P., Rusconi, F., Cañón, P., Medina, C., Conti, L., Cominelli, E., Tonelli, C., & Arce-Johnson, P. (2011). The grapevine guard cell-related VvMYB60 transcription factor is involved in the regulation of stomatal activity and is differentially expressed in response to ABA and osmotic stress. BMC Plant Biology, 11(1). 10.1186/1471-2229-11-142

Ghan, R., Van Sluyter, S. C., Hochberg, U., Degu, A., Hopper, D. W., Tillet, R. L., Schlauch, K. A., et al. (2015, November 16). Five omic technologies are concordant in differentiating the biochemical characteristics of the berries of five grapevine (Vitis vinifera L.) cultivars. BMC Genomics. Springer Science and Business Media LLC. Retrieved from 10.1186/s12864-015-2115-y

Giacomelli, L., Zeilmaker, T., Scintilla, S., Salvagnin, U., van der Voort, J. R., & Moser, C. (2022, April 19). Vitis vinifera plants edited in DMR6 genes show improved resistance to downy mildew. Cold Spring Harbor Laboratory. Retrieved from 10.1101/2022.04.19.488768

Giacomelli, L., Zeilmaker, T., Giovannini, O., Salvagnin, U., Masuero, D., Franceschi, P., Vrhovsek, U., et al. (2023, August 21). Simultaneous editing of two DMR6 genes in grapevine results in reduced susceptibility to downy mildew. Frontiers in Plant Science. Frontiers Media SA. Retrieved from 10.3389/fpls.2023.1242240

Grimplet, J., Pimentel, D., Agudelo-Romero, P., Martinez-Zapater, J. M., & Fortes, A. M. (2017). The LATERAL ORGAN BOUNDARIES Domain gene family in grapevine: genome-wide characterization and expression analyses during developmental processes and stress responses. Scientific Reports, 7(1). 10.1038/s41598-017-16240-5

Gupta, A., Rico-Medina, A., & Caño-Delgado, A. I. (2020, April 17). The physiology of plant responses to drought. Science. American Association for the Advancement of Science (AAAS). Retrieved from 10.1126/science.aaz7614

Hewitt, S., Hernández-Montes, E., Dhingra, A., & Keller, M. (2023). Impact of heat stress, water stress, and their combined effects on the metabolism and transcriptome of grape berries. Scientific Reports, 13(1). 10.1038/s41598-023-36160-x

Hochberg, U., Rockwell, F. E., Holbrook, N. M., & Cochard, H. (2018, February). Iso/Anisohydry: A Plant–Environment Interaction Rather Than a Simple Hydraulic Trait. Trends in Plant Science. Elsevier BV. Retrieved from 10.1016/j.tplants.2017.11.002

Hunt, L., Bailey, K. J., & Gray, J. E. (2010). The signalling peptide EPFL9 is a positive regulator of stomatal development. New Phytologist, 186(3), 609–614. 10.1111/j.1469-8137.2010.03200.x

IMARC; Wine Market Size, Share | Global Industry Trends 2032. https://www.imarcgroup.com/wine-market

Imes, D., Mumm, P., Böhm, J., Al-Rasheid, K. A. S., Marten, I., Geiger, D., & Hedrich, R. (2013). Open stomata 1 (OST1) kinase controls R–type anion channel QUAC1 in Arabidopsis guard cells. The Plant Journal, 74(3), 372–382. 10.1111/tpj.12133

Ilc, T., Arista, G., Tavares, R., Navrot, N., Duchêne, E., Velt, A., Choulet, F., Paux, E., Fischer, M., Nelson, D. R., Hugueney, P., Werck-Reichhart, D., & Rustenholz, C. (2018). Annotation, classification, genomic organization and expression of the Vitis vinifera CYPome. PLoS ONE, 13(6), e0199902. 10.1371/journal.pone.0199902

Jaldhani, V., Sanjeeva Rao, D., Beulah, P., Nagaraju, P., Suneetha, K., Veronica, N., Kondamudi, R., et al. (2022). Drought and heat stress combination in a changing climate. Climate Change and Crop Stress. Elsevier. Retrieved from 10.1016/B978-0-12-816091-6.00002-X

Khadka, V. S., Vaughn, K., Xie, J., Swaminathan, P., Ma, Q., Cramer, G. R., & Fennell, A. Y. (2019). Transcriptomic response is more sensitive to water deficit in shoots than roots of Vitis riparia (Michx.). BMC Plant Biology, 19(1). 10.1186/s12870-019-1664-7

Konecny, T., Asatryan, A., Nikoghosyan, M., & Binder, H. (2024, September 6). Unveiling Iso- and Aniso-Hydric Disparities in Grapevine—A Reanalysis by Transcriptome Portrayal Machine Learning. Plants. MDPI AG. Retrieved from 10.3390/plants13172501

Lau, O. S., & Bergmann, D. C. (2012). Stomatal development: a plant’s perspective on cell polarity, cell fate transitions and intercellular communication. Development, 139(20), 3683–3692. 10.1242/dev.080523

Lamarque, L. J., Delmas, C. E. L., Charrier, G., Burlett, R., Dell’Acqua, N., Pouzoulet, J., Gambetta, G. A., et al. (2023, May 12). Quantifying the grapevine xylem embolism resistance spectrum to identify varieties and regions at risk in a future dry climate. Scientific Reports. Springer Science and Business Media LLC. Retrieved from 10.1038/s41598-023-34224-6

Le, J., Zou, J., Yang, K., & Wang, M. (2014). Signaling to stomatal initiation and cell division. Frontiers In Plant Science, 5. 10.3389/fpls.2014.00297

Leng, X., Wang, P., Wang, C., Zhu, X., Li, X., Li, H., Mu, Q., Li, A., Liu, Z., & Fang, J. (2017). Genome-wide identification and characterization of genes involved in carotenoid metabolic in three stages of grapevine fruit development. Scientific Reports, 7(1). 10.1038/ s41598-017-04004-0

Li, D., Gu, B., Huang, C., Shen, J., Wang, X., Guo, J., Yu, R., Mou, S., & Guan, Q. (2023). Functional Study of Amorpha fruticosa WRKY20 Gene in Response to Drought Stress. International Journal Of Molecular Sciences, 24(15), 12231. 10.3390/ijms241512231

Li, M., Duan, Z., Zhang, S., Zhang, J., Chen, J., & Song, H. (2024). The physiological and molecular mechanisms of WRKY transcription factors regulating drought tolerance: A review. Gene, 938, 149176. 10.1016/j.gene.2024.149176

Liao, Y., Smyth, G. K., & Shi, W. (2013, November 13). featureCounts: an efficient general purpose program for assigning sequence reads to genomic features. Bioinformatics. Oxford University Press (OUP). Retrieved from 10.1093/bioinformatics/btt656

Lin, Y., Liu, S., Fang, X., Ren, Y., You, Z., Xia, J., Hakeem, A., Yang, Y., Wang, L., Fang, J., & Shangguan, L. (2023). The physiology of drought stress in two grapevine cultivars: Photosynthesis, antioxidant system, and osmotic regulation responses. Physiologia Plantarum, 175(5). 10.1111/ppl.14005

Liu, J., Zhao, F., Guo, Y., Fan, X., Wang, Y., & Wen, Y. (2019). The ABA receptor-like gene VyPYL9 from drought-resistance wild grapevine confers drought tolerance and ABA hypersensitivity in Arabidopsis. Plant Cell Tissue And Organ Culture (PCTOC), 138(3), 543–558. 10.1007/s11240-019-01650-2

Love, M. I., Huber, W., & Anders, S. (2014, December 5). Moderated estimation of fold change and dispersion for RNA-seq data with DESeq2. Genome Biology. Springer Science and Business Media LLC. Retrieved from 10.1186/s13059-014-0550-8

Luo, X., Bai, X., Sun, X., Zhu, D., Liu, B., Ji, W., Cai, H., Cao, L., Wu, J., Hu, M., Liu, X., Tang, L., & Zhu, Y. (2013). Expression of wild soybean WRKY20 in Arabidopsis enhances drought tolerance and regulates ABA signalling. Journal Of Experimental Botany, 64(8), 2155–2169. 10.1093/jxb/ert073

Ma, Q., & Yang, J. (2018). Transcriptome profiling and identification of functional genes involved in H2S response in grapevine tissue cultured plantlets. Genes & Genomics, 40(12), 1287–1300. 10.1007/s13258-018-0723-z

Magon, G., De Rosa, V., Martina, M., Falchi, R., Acquadro, A., Barcaccia, G., Portis, E., et al. (2023, December 11). Boosting grapevine breeding for climate-smart viticulture: from genetic resources to predictive genomics. Frontiers in Plant Science. Frontiers Media SA. Retrieved from 10.3389/fpls.2023.1293186

Malnoy, M., Viola, R., Jung, M.-H., Koo, O.-J., Kim, S., Kim, J.-S., Velasco, R., et al. (2016, December 20). DNA-Free Genetically Edited Grapevine and Apple Protoplast Using CRISPR/Cas9 Ribonucleoproteins. Frontiers in Plant Science. Frontiers Media SA. Retrieved from 10.3389/fpls.2016.01904

Meggio, F., Prinsi, B., Negri, A. S., Simone Di Lorenzo, G., Lucchini, G., Pitacco, A., Failla, O., et al. (2014, March 20). Biochemical and physiological responses of two grapevine rootstock genotypes to drought and salt treatments. Australian Journal of Grape and Wine Research. Hindawi Limited. Retrieved from 10.1111/ajgw.12071

Mishra, A. K., & Singh, V. P. (2010, September). A review of drought concepts. Journal of Hydrology. Elsevier BV. Retrieved from 10.1016/j.jhydrol.2010.07.012

Moretto, M., Sonego, P., Pilati, S., Malacarne, G., Costantini, L., Grzeskowiak, L., Bagagli, G., et al. (2016, May 10). VESPUCCI: Exploring Patterns of Gene Expression in Grapevine. Frontiers in Plant Science. Frontiers Media SA. Retrieved from 10.3389/fpls.2016.00633

Moretto, M., Sonego, P., Pilati, S., Matus, J. T., Costantini, L., Malacarne, G., & Engelen, K. (2022, February 24). A COMPASS for VESPUCCI: A FAIR Way to Explore the Grapevine Transcriptomic Landscape. Frontiers in Plant Science. Frontiers Media SA. Retrieved from 10.3389/fpls.2022.815443

Navarro-Payá, D., Santiago, A., Orduña, L., Zhang, C., Amato, A., D’Inca, E., Fattorini, C., et al. (2022, January 17). The Grape Gene Reference Catalogue as a Standard Resource for Gene Selection and Genetic Improvement. Frontiers in Plant Science. Frontiers Media SA. Retrieved from 10.3389/fpls.2021.803977

Nerva, L., Chitarra, W., Fila, G., Lovat, L., & Gaiotti, F. (2023). Variability in Stomatal Adaptation to Drought among Grapevine Cultivars: Genotype-Dependent Responses. Agriculture, 13(12), 2186. 10.3390/agriculture13122186

Nicolas, P., Lecourieux, D., Kappel, C., Cluzet, S., Cramer, G., Delrot, S., & Lecourieux, F. (2013). The Basic Leucine Zipper Transcription Factor ABSCISIC ACID RESPONSE ELEMENT-BINDING FACTOR2 Is an Important Transcriptional Regulator of Abscisic Acid-Dependent Grape Berry Ripening Processes. PLANT PHYSIOLOGY, 164(1), 365–383. 10.1104/pp.113.231977

Orduña, L., Li, M., Navarro-Payá, D., Zhang, C., Santiago, A., Romero, P., Ramšak, Ž., et al. (2022, March 3). Direct regulation of shikimate, early phenylpropanoid, and stilbenoid pathways by Subgroup 2 R2R3-MYBs in grapevine. The Plant Journal. Wiley. Retrieved from 10.1111/tpj.15686

Orduña, L., Santiago, A., Navarro-Payá, D., Zhang, C., Wong, D. C. J., & Matus, J. T. (2023, September 5). Aggregated gene co-expression networks predict transcription factor regulatory landscapes in grapevine. (S. Osorio, Ed.)Journal of Experimental Botany. Oxford University Press (OUP). Retrieved from 10.1093/jxb/erad344

Pessina, S., Lenzi, L., Perazzolli, M., Campa, M., Dalla Costa, L., Urso, S., Valè, G., et al. (2016, April 20). Knockdown of MLO genes reduces susceptibility to powdery mildew in grapevine. Horticulture Research. Oxford University Press (OUP). Retrieved from 10.1038/hortres.2016.16

Pilati, S., Malacarne, G., Navarro-Payá, D., Tomè, G., Riscica, L., Cavecchia, V., Matus, J. T., et al. (2021, November 23). Vitis OneGenE: A Causality-Based Approach to Generate Gene Networks in Vitis vinifera Sheds Light on the Laccase and Dirigent Gene Families. Biomolecules. MDPI AG. Retrieved from 10.3390/biom11121744

Pilati, S., Bagagli, G., Sonego, P., Moretto, M., Brazzale, D., Castorina, G., Simoni, L., Tonelli, C., Guella, G., Engelen, K., Galbiati, M., & Moser, C. (2017). Abscisic Acid Is a Major Regulator of Grape Berry Ripening Onset: New Insights into ABA Signaling Network. Frontiers In Plant Science, 8. 10.3389/fpls.2017.01093

Pirrello, C., Malacarne, G., Moretto, M., Lenzi, L., Perazzolli, M., Zeilmaker, T., Van den Ackerveken, G., et al. (2022, January 22). Grapevine DMR6-1 Is a Candidate Gene for Susceptibility to Downy Mildew. Biomolecules. MDPI AG. Retrieved from 10.3390/biom12020182

Ritchie, M. E., Phipson, B., Wu, D., Hu, Y., Law, C. W., Shi, W., & Smyth, G. K. (2015, January 20). limma powers differential expression analyses for RNA-sequencing and microarray studies. Nucleic Acids Research. Oxford University Press (OUP). Retrieved from 10.1093/nar/gkv007

Rodriguez-Izquierdo, A., Carrasco, D., Valledor, L., Bota, J., López-Hidalgo, C., Revilla, M. A., & Arroyo-Garcia, R. (2024). The scion-driven transcriptomic changes guide the resilience of grafted near-isohydric grapevines under water deficit. Horticulture Research, 12(2). 10.1093/hr/uhae291

Roka, K. (2019). Anthropocene and Climate Change. Encyclopedia of the UN Sustainable Development Goals. Springer International Publishing. Retrieved from 10.1007/978-3-319-71063-1_26-1

Rossdeutsch, L., Edwards, E., Cookson, S. J., Barrieu, F., Gambetta, G. A., Delrot, S., & Ollat, N. (2016). ABA-mediated responses to water deficit separate grapevine genotypes by their genetic background. BMC Plant Biology, 16(1). 10.1186/s12870-016-0778-4

Saito, S., Hirai, N., Matsumoto, C., Ohigashi, H., Ohta, D., Sakata, K., & Mizutani, M. (2004). Arabidopsis CYP707As Encode (+)-Abscisic Acid 8′-Hydroxylase, a Key Enzyme in the Oxidative Catabolism of Abscisic Acid. PLANT PHYSIOLOGY, 134(4), 1439–1449. 10.1104/pp.103.037614

Santiago, A., Orduña, L., Fernández, J. D., Vidal, Á., De Martín-Agirre, I., Lisón, P., Vidal, E. A., Navarro-Payá, D., & Matus, J. T. (2024). The Plantae Visualization Platform: a comprehensive web-based tool for the integration, visualization, and analysis of omic data across plant and related species. bioRxiv (Cold Spring Harbor Laboratory). 10.1101/2024.12.19.629382

Savoi, S., Wong, D. C. J., Arapitsas, P., Miculan, M., Bucchetti, B., Peterlunger, E., Fait, A., et al. (2016, March 21). Transcriptome and metabolite profiling reveals that prolonged drought modulates the phenylpropanoid and terpenoid pathway in white grapes (Vitis vinifera L.). BMC Plant Biology. Springer Science and Business Media LLC. Retrieved from 10.1186/s12870-016-0760-1

Savoi, S., Wong, D. C. J., Degu, A., Herrera, J. C., Bucchetti, B., Peterlunger, E., Fait, A., Mattivi, F., & Castellarin, S. D. (2017). Multi-Omics and Integrated Network Analyses Reveal New Insights into the Systems Relationships between Metabolites, Structural Genes, and Transcriptional Regulators in Developing Grape Berries (Vitis vinifera L.) Exposed to Water Deficit. Frontiers In Plant Science, 8. 10.3389/fpls.2017.01124

Seo, P. J., Xiang, F., Qiao, M., Park, J., Lee, Y. N., Kim, S., Lee, Y., Park, W. J., & Park, C. (2009). The MYB96 Transcription Factor Mediates Abscisic Acid Signaling during Drought Stress Response in Arabidopsis. PLANT PHYSIOLOGY, 151(1), 275–289. 10.1104/pp.109.144220

Singh, B. K., Delgado-Baquerizo, M., Egidi, E., Guirado, E., Leach, J. E., Liu, H., & Trivedi, P. (2023, May 2). Climate change impacts on plant pathogens, food security and paths forward. Nature Reviews Microbiology. Springer Science and Business Media LLC. Retrieved from 10.1038/s41579-023-00900-7

Tan, J. W., Shinde, H., Tesfamicael, K., Hu, Y., Fruzangohar, M., Tricker, P., Baumann, U., Edwards, E. J., & López, C. M. R. (2023). Global transcriptome and gene co-expression network analyses reveal regulatory and non-additive effects of drought and heat stress in grapevine. Frontiers In Plant Science, 14. 10.3389/fpls.2023.1096225

Tang, L., Cai, H., Zhai, H., Luo, X., Wang, Z., Cui, L., & Bai, X. (2014). Overexpression of Glycine soja WRKY20 enhances both drought and salt tolerance in transgenic alfalfa (Medicago sativa L.). Plant Cell Tissue And Organ Culture (PCTOC), 118(1), 77–86. 10.1007/s11240-014-0463-y

Tang, X., Cao, X., Xu, X., Jiang, Y., Luo, Y., Ma, Z., Fan, J., et al. (2017, October). Effects of Climate Change on Epidemics of Powdery Mildew in Winter Wheat in China. Plant Disease. Scientific Societies. Retrieved from 10.1094/PDIS-02-17-0168-RE

Trenti, M., Lorenzi, S., Bianchedi, P. L., Grossi, D., Failla, O., Grando, M. S., & Emanuelli, F. (2021, January 6). Candidate genes and SNPs associated with stomatal conductance under drought stress in Vitis. BMC Plant Biology. Springer Science and Business Media LLC. Retrieved from 10.1186/s12870-020-02739-z

Tu, M., Wang, X., Zhu, Y., Wang, D., Zhang, X., Cui, Y., Li, Y., Gao, M., Li, Z., Wang, Y., & Wang, X. (2018). VlbZIP30 of grapevine functions in dehydration tolerance via the abscisic acid core signaling pathway. Horticulture Research, 5(1). 10.1038/s41438-018-0054-x

Vitulo, N., Forcato, C., Carpinelli, E. C., Telatin, A., Campagna, D., D’Angelo, M., Zimbello, R., et al. (2014, April 17). A deep survey of alternative splicing in grape reveals changes in the splicing machinery related to tissue, stress condition and genotype. BMC Plant Biology. Springer Science and Business Media LLC. Retrieved from 10.1186/1471-2229-14-99

Van Leeuwen, C., Sgubin, G., Bois, B., Ollat, N., Swingedouw, D., Zito, S., & Gambetta, G. A. (2024, March 26). Climate change impacts and adaptations of wine production. Nature Reviews Earth & Environment. Springer Science and Business Media LLC. Retrieved from 10.1038/s43017-024-00521-5

Vannozzi, A., Palumbo, F., Magon, G., Lucchin, M., & Barcaccia, G. (2021, September 1). The grapevine (Vitis vinifera L.) floral transcriptome in Pinot noir variety: identification of tissue-related gene networks and whorl-specific markers in pre-and post-anthesis phases. Horticulture Research. Oxford University Press (OUP). Retrieved from 10.1038/s41438-021-00635-7

Wang, Y., Zhang, R., Liang, Z., & Li, S. (2020, March 16). Grape-RNA: A Database for the Collection, Evaluation, Treatment, and Data Sharing of Grape RNA-Seq Datasets. Genes. MDPI AG. Retrieved from 10.3390/genes11030315

Wang, S., Zhou, Z., Rahiman, R., Lee, G. S. Y., Yeo, Y. K., Yang, X., & Lau, O. S. (2021). Light regulates stomatal development by modulating paracrine signaling from inner tissues. Nature Communications, 12(1). 10.1038/s41467-021-23728-2

Waseem, M., Nkurikiyimfura, O., Niyitanga, S., Jakada, B. H., Shaheen, I., & Aslam, M. M. (2022). GRAS transcription factors emerging regulator in plants growth, development, and multiple stresses. Molecular Biology Reports, 49(10), 9673–9685. 10.1007/s11033-022-07425-x

Winterhagen, P., Howard, S. F., Qiu, W., & Kovács, L. G. (2008, June). Transcriptional Up-Regulation of GrapevineMLOGenes in Response to Powdery Mildew Infection. American Journal of Enology and Viticulture. American Society for Enology and Viticulture. Retrieved from 10.5344/ajev.2008.59.2.159

Wong, D. C. J., Schlechter, R., Vannozzi, A., Höll, J., Hmmam, I., Bogs, J., Tornielli, G. B., et al. (2016, July 12). A systems-oriented analysis of the grapevine R2R3-MYB transcription factor family uncovers new insights into the regulation of stilbene accumulation. DNA Research. Oxford University Press (OUP). Retrieved from 10.1093/dnares/dsw028

Wong, D. C. J. (2020, April 7). Network aggregation improves gene function prediction of grapevine gene co-expression networks. Plant Molecular Biology. Springer Science and Business Media LLC. Retrieved from 10.1007/s11103-020-01001-2

Zhang, R., Wang, Y., Li, S., Yang, L., & Liang, Z. (2020). ABA signaling pathway genes and function during abiotic stress and berry ripening in Vitis vinifera. Gene, 769, 145226. 10.1016/j.gene.2020.14522

Zhao, H., Xu, D., Tian, T., Kong, F., Lin, K., Gan, S., Zhang, H., & Li, G. (2020). Molecular and functional dissection of EARLY-FLOWERING 3 (ELF3) and ELF4 in Arabidopsis. Plant Science, 303, 110786. 10.1016/j.plantsci.2020.110786

